# The neural dynamics associated with computational complexity

**DOI:** 10.1101/2022.01.05.475102

**Authors:** Juan Pablo Franco, Peter Bossaerts, Carsten Murawski

**Affiliations:** Brain, Mind & Markets Laboratory, Department of Finance, The University of Melbourne, Melbourne, Victoria, Australia; Florey Institute of Neuroscience and Mental Health, Melbourne, Victoria, Australia

**Keywords:** decision-making, computational complexity, typical-case complexity, knapsack problem, problem-solving, cognitive control, anterior cingulate cortex (ACC), anterior insula (AI), fMRI

## Abstract

Many everyday tasks require people to solve computationally complex problems. However, little is known about the effects of computational hardness on the neural processes associated with solving such problems. Here, we draw on computational complexity theory to address this issue. We performed an experiment in which participants solved several instances of the 0-1 knapsack problem, a combinatorial optimization problem, while undergoing ultra-high field (7T) functional magnetic resonance imaging (fMRI). Instances varied in two task-independent measures of intrinsic computational hardness: complexity and proof hardness. We characterise a network of brain regions whose activation was correlated with both measures but in distinct ways, including the anterior insula, dorsal anterior cingulate cortex and the intra-parietal sulcus/angular gyrus. Activation and connectivity changed dynamically as a function of complexity and proof hardness, in line with theoretical computational requirements. Overall, our results suggest that computational complexity theory provides a suitable framework to study the effects of computational hardness on the neural processes associated with solving complex cognitive tasks.

## 1. Introduction

Every day, people make decisions that require solving complex problems. Many of these problems are known to be computationally intractable in the sense that the number of operations that need to be performed to find a solution grows quickly to levels that makes solving correctly these problems infeasible. Real-life examples of intractable tasks include attention gating, task scheduling, shopping, routing, bin packing, and game play (van Rooij et al., 2019; Bossaerts and Murawski, 2017). Despite the relevance of intractable problems in daily life, little is known about the effects of complexity of tasks on the neural processes during problem-solving.

Intractable problems require an extended period of time to solve and involve an extensive search space. These two characteristics defy formal investigation of neural dynamics. Firstly, since solving such tasks requires an extended period of time, one cannot opt for modeling based on discrete choice theories such as those underlying neuroeconomics (Yoo et al., 2021). When deciding between, say, an apple and a candy, the neural activation can be modeled in terms of an indicator variable whose level is modulated by the value inferred from choices and kept constant throughout the short (couple of seconds) deliberation time (e.g., Hare et al., 2011). When choice concerns complex alternatives, deliberation times may be an order of magnitude longer, so neural activation can be expected to fluctuate markedly during the course of deliberation. Secondly, because the search space is large, there are a plethora of paths that can be chosen during resolution. Since human approaches to solving a complex problem exhibit substantial heterogeneity, both across individuals and over time (e.g., Murawski and Bossaerts, 2016), modeling neural dynamics during deliberation is bound to be challenging if it is to be based on “what people are thinking,” i.e., on individual approaches to solving complex problems. A different strategy is called for.

Here, we propose to focus on “what people are solving,” that is, on features of the computational tasks that are being presented. This has precedent in the analysis of probabilistic tasks, where intrinsic properties of the gamble at hand (such as mean and variance of the uncertain reward) have proven invaluable to deciphering the neural processes leading up to choice (e.g., D’Acremont and Bossaerts, 2008). Likewise, mathematical characteristics of the stimuli in perceptual tasks, such as signal strength, elucidate neural dynamics during deliberation (e.g., Ploran et al., 2011; Hanks and Summerfield, 2017). Drawing on computational complexity theory, we demonstrate here that a mapping exists between intrinsic properties of instances of a problem related to computational hardness and neural dynamics during decision-making. These properties represent generic features of computational problems that can be studied across different tasks.

We studied the case of a canonical NP-complete (i.e., both NP-hard and NP) problem, the 0-1 knapsack decision problem (KP). There, the decision-maker is asked to choose whether, given a set of items with differing value and weight, there exists a subset whose total value is at least as a high as a given threshold, while the total weight is less than or equal to a capacity constraint. We identified two properties of instances of KP related to an instance’s computational hardness and tested whether these properties elucidated neural signatures during deliberation. The two properties are *complexity* and *proof hardness*. Complexity captures the number of computational steps (or time) needed to solve an instance, while proof hardness represents the computational steps needed to verify the correctness of the solution.

In order to measure complexity, we utilized a metric of difficulty that arises from the study of *random ensembles of instances* (i.e., random cases of the problem). Variation in expected computational complexity, regardless of the algorithm used, has been attributed to specific structural properties of instances (Cheeseman et al., 1991; Percus et al., 2006; Gent et al., 1996; Yadav et al., 2020; Achlioptas et al., 2005; Selman and Kirkpatrick, 1996; Krzakala et al., 2006). The resulting “typical-case complexity” (TCC) has been found to affect human performance and effort in several intractable (NP-hard) problem-solving tasks, including KP (Franco et al., 2021a,b). Therefore, we hypothesized that TCC would prove useful in characterising the effects of computational hardness on neural processes. In analogy with work on neural correlates related to deliberation during tractable tasks (Fedorenko et al., 2013; Assem et al., 2020; Duncan and Owen, 2000; Crittenden et al., 2016), we expected neural correlates of TCC to overlap with the mutliple-demand system (MDS). Specifically, we hypothesized they would overlap with two networks, (1) the cingulo-opercular network (CON), consisting of the dorsal anterior cingulate cortex (dACC) and the anterior insula (AI), and (2) the frontoparietal network (FPN), composed of the intraparietal sulcus (IPS) and specific regions from the lateral prefrontal cortex including the inferior frontal sulcus and the middle frontal gyrus (MFG) (e.g., Crittenden et al. (2016); Fedorenko et al. (2013); Duncan (2010); Dosenbach et al. (2007)). Additionally, we expected the level of complexity to be associated with neural markers of efficacy (Fromer et al., 2021) and performance (Neta et al., 2017; Bossaerts, 2018).

We appeal to the theory of proof complexity to measure proof hardness. In the context of an NP-complete problem, such as the KP, there exists an asymmetry in the difficulty of proving that the solution is correct, which depends on the “satisfiability” of the instance. If an instance is *satisfiable* (the correct choice is ‘yes’), it suffices to find a witness (example assignment of variables) that satisfies all of the constraints; one can then quickly verify that the witness indeed satisfies all the constraints, and this verification can be done in polynomial time. To confirm that an instance is *unsatisfiable* (the correct choice is ‘no’) requires proving that *no* witness exists, which is far more difficult: even if a few potential witnesses are found not to be feasible, there may exist others that are. Theoretically, the asymmetry reflects the conjectured null intersection between complexity classes NP-complete and co-NP-complete (Arora and Barak, 2009). We thus conjectured that satisfiability would correlate with subjective markers of *reliability*, that is, the degree to which the result of a calculation can be relied on to be accurate—much like variance modulates subjective beliefs of choice correctness in probabilistic tasks. Therefore, we expected neural correlates of this measure in regions that have been previously shown to encode uncertainty, specifically in CON (Neta et al., 2017, 2014; Bossaerts, 2018; Fouragnan et al., 2018).

In our experiment, participants were asked to solve several instances of the knapsack decision problem, while undergoing functional magnetic resonance imaging (fMRI). Instances were drawn randomly but in a way that systematically varied their TCC and their satisfiability. Critically, in order to more precisely localize and track neural signals during deliberation, we employed ultra-high field (7 Tesla) fMRI.

## 2. Results

Twenty participants (14 female, 5 male, 1 other; age range = 18-35 years, mean age = 26.6 years) took part in this study. Each participant was asked to solve 56 instances of the knapsack decision task while undergoing ultra-high field MRI brain scanning (Fig 1). Instances varied in their computational complexity (TCC) and their satisfiability (2*×*2 balanced factorial design; see Materials and Methods).

**Figure 1:**
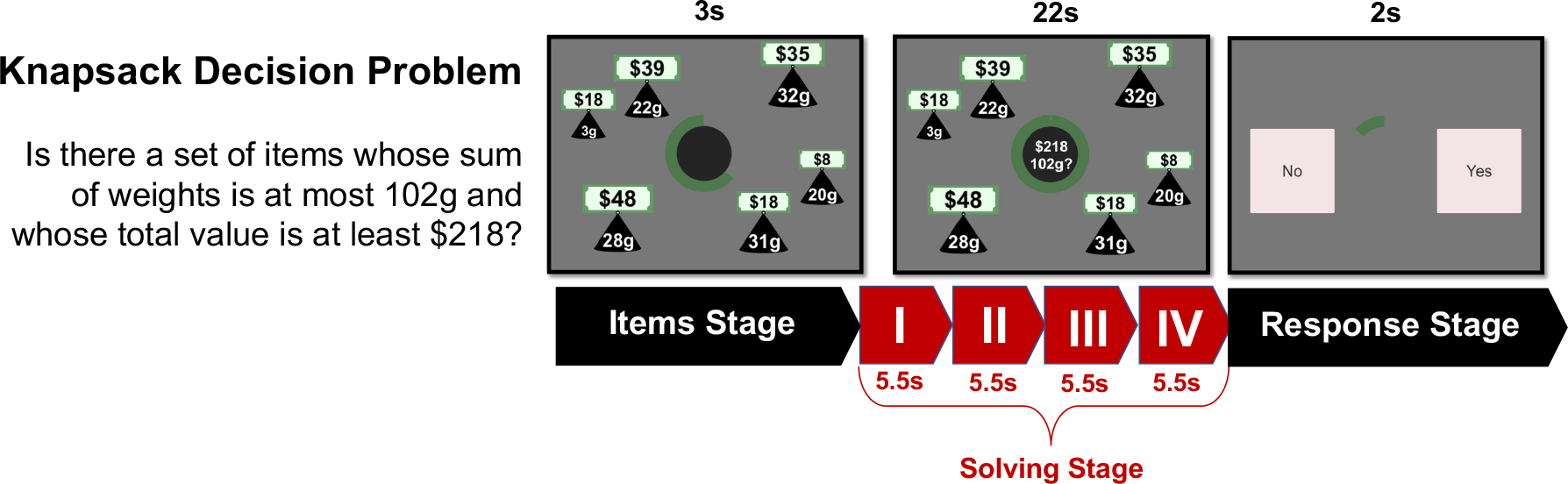
Knapsack decision task. The task was composed of three main stages: items stage (3s), solving stage (22s) and response stage (2s). Initially, participants were presented with a set of items of different values and weights. The green circle at the center of the screen indicated the time remaining in this stage of the trial. This stage lasted 3 seconds. Then, both capacity constraint and target profit were shown at the center of the screen. The objective of the task is to decide whether there exists a subset of items for which (1) the sum of weights is lower or equal to the capacity constraint and (2) the sum of values yields at least the target profit. This stage lasted 22 seconds. Finally, participants had 2 seconds to make either a ‘YES’ or ‘NO’ response using the keyboard. A fixation cross was shown during the inter-trial interval (jittered between 8 and 12 seconds). Initially, for the neuroimaging data analysis the solving stage was partitioned into four Boxcar response functions (periods S1-S4), while the items and response stage were modeled with a single Boxcar function for the duration of the stage.

Additionally, participants performed, outside the scanner, a set of complementary tasks, including a knapsack optimization task and a set of cognitive function tasks. In this section, we report the behavioral results of the knapsack decision task, while the behavioral results from the complementary tasks are reported in Appendices D and C.2.

### 2.1. Behavioral results

#### 2.1.1. Summary statistics

On average, participants chose the ‘YES’ option 50% of the trials (min = 25%, max = 68%). Mean *human performance*, measured as the proportion of trials in which a correct response was made, was 0.78 (min = 0.48, max = 0.95, *SD* = 0.14). Performance increased slightly as the task progressed but the change was not statistically significant (*β*_0.5_ = 0.009, *HDI*_0.95_ = [*−*0.001, 0.021], main effect of trial number on performance, generalized logistic mixed model (GLMM); Table 4 Model 1).

#### 2.1.2. Accuracy and instance properties

We first studied the effect of TCC on human performance. This measure is based on a prominent framework in computer science that investigates the factors affecting computational hardness in computational problems by studying the difficulty of randomly generated instances of those problems. In the knapsack problem, TCC is explicitly connected to a set of parameters 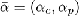 that capture the constrainedness of the problem: 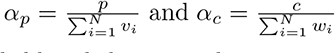 (Yadav et al., 2020; Franco et al., 2021b). These parameters determine the likelihood that a random instance is *satisfiable*, that is, that the solution is ‘yes’. Specifically, they characterize where typical instances are generally satisfiable (under-constrained region), where they are unsatisfiable (over-constrained region) and where the probability of satisfiability is close to 50% (satisfiability threshold *α_s_*). It has been shown that the computational difficulty of solving the problem is higher when *α_p_* is close to *α_s_* (Yadav et al., 2020; Franco et al., 2021b). TCC is then defined based on the distance of *α_p_* to the satisfiability threshold *α_s_*. Specifically, instances with values of *α_p_* near the satisfiability threshold have a high typical-case complexity (*high TCC* ) whereas instances further away from it—that is, in the under-constrained and over-constrained regions— have low typical-case complexity (*low TCC* ). In line with previous results (Franco et al., 2021b), we found participants had a better performance on instances with low TCC compared to those with high TCC (*β*_0.5_ = *−*1.10, *HDI*_0.95_ = [*−*1.44*, −*0.79], main effect of TCC on performance, GLMM; Fig 2; Table 4 Model 2).

**Figure 2:**
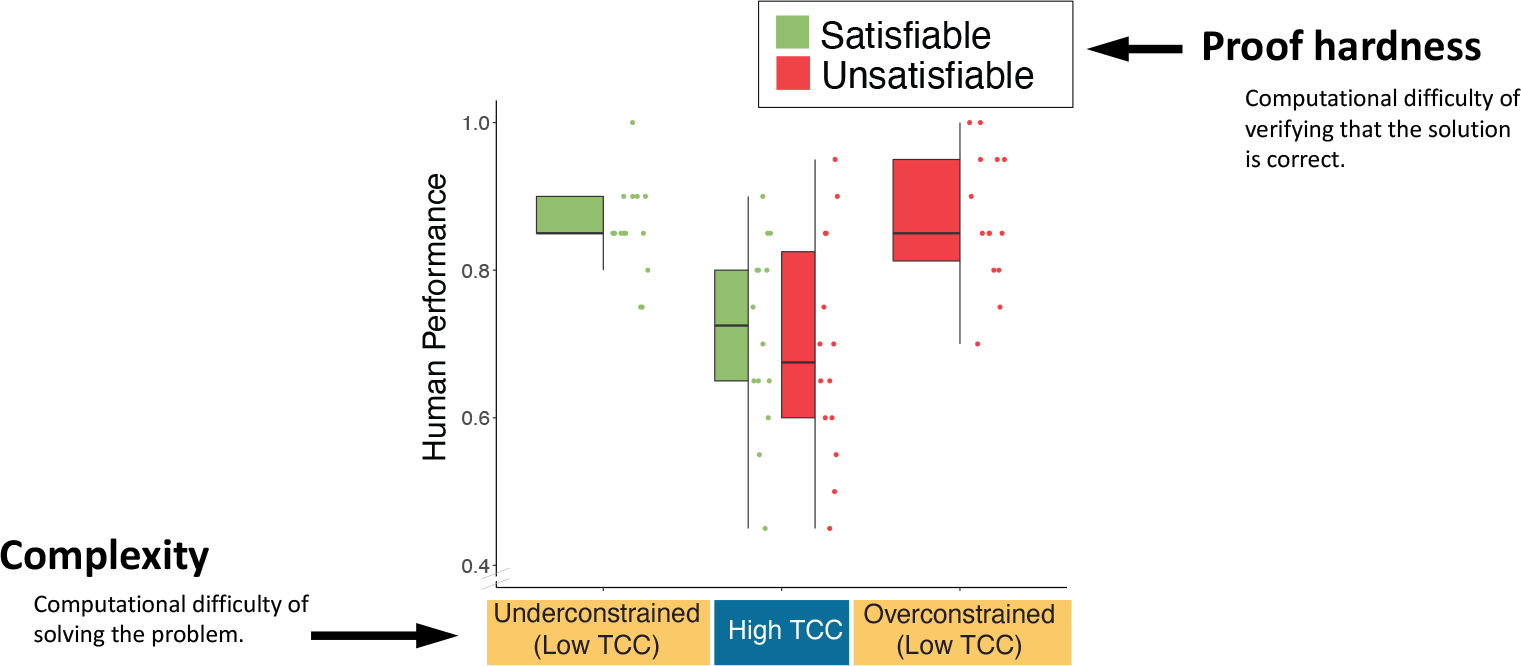
Relation between TCC and human performance in the knapsack decision task. Each dot represents an instance; human performance corresponds the proportion of participants that solved the instance correctly. Instances are categorized according to their constrainedness region (*α*) and their TCC. In the underconstrained region (low TCC) the satisfiability probability is close to one, while in the overconstrained region (low TCC) the probability is close to zero. The region with a high TCC corresponds to a region in which the probability is close to 0.5. Additionally, instances are categorized according to their solution (satisfiability) which is represented by their color. *The box-plots represent the median, the interquartile range (IQR) and the whiskers extend to a maximum length of 1.5*IQR*

We then studied satisfiability. This property of an instance captures the computational difficulty of verifying that the certificate of a solutions (i.e., proof) is correct. To verify that an instance is satisfiable, it suffices to check that a candidate set of items (satisfiability-certificate) satisfies the constraints. In contrast, verifying unsatisfiability requires validating a proof of non-existence (unsatisfiability-certificate). For NP-complete problems the former is tractable (P-time) whilst the latter is conjectured to be intractable (follows from the conjecture that *coNP /*= *NP* ; Arora and Barak (2009)). Our findings replicate previous results that suggest there is no effect of satisfiability on human performance in the knapsack decision task (Franco et al. (2021b); *β*_0.5_ = 0.02, *HDI*_0.95_ = [*−*0.30, 0.30], main effect of satisfiability on performance, GLMM; Table 4 Model 5). Moreover, we found no significant interaction effect between TCC and satisfiability on performance (*β*_0.5_ = 0.26, *HDI*_0.95_ = [*−*0.37, 0.90], interaction effect of TCC and satisfiability, GLMM; Fig 2; Table 4 Model 6). Besides studying and replicating previously reported effects of TCC and satisfiability on human performance, we replicated other key findings presented by Franco et al. (2021b) (Appendix C).

Finally, we investigated human performance in a set of related tasks. We explored the relation between performance in the knapsack tasks and core cognitive abilities, including working memory, episodic memory, strategy use, as well as mental arithmetic. For this analysis we utilized the joined data set from this study together with data collected by Franco et al. (2021b). Our results suggest a weak relation between these cognitive abilities and performance in the knapsack tasks. The only significant correlation (at *α* = 0.05) shows a link between mental arithmetic ability and performance in the knapsack optimization task (Appendix D).

### 2.2. Imaging results

#### 2.2.1. Whole-brain analysis

We conducted a whole-brain analysis of the neural correlates of two intrinsic generic properties of problems: TCC and satisfiability. Additionally, we investigated the neural correlates of response accuracy (Appendix E). We did this by fitting GLMs that partitioned the solving stage into four separate periods *Neural correlates of TCC.* We expected to see the *highTCC − lowTCC* contrast capture differences in BOLD activation in regions previously correlated with cognitive demand (i.e., MDS). We explicitly expected to find evidence for the encoding of TCC in CON from early on during the solving stage due to its link with expected performance and reliability. Higher TCC entails, on average, lower performance and lower reliability of finding the solution (Fig 2). Note that the estimation of TCC early on in the trial is feasible, because constrainedness (and thus TCC) can be potentially estimated by performing sum and division operations 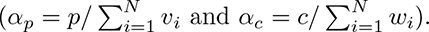.

We found that the neural correlates of TCC varied throughout the duration of the solving stage (Fig 3, Table 1). Contrary to our expectations, we did not find significant correlations of TCC during the first period of the solving stage. Interestingly, during the second period we did find a set of clusters that showed higher BOLD activity on instances with low TCC. These regions include the angular gyrus (AG) bilaterally, the SFG, the right MFG as well as regions in the orbitofrontal cortex (bilaterally). It is worth noting that the negative pattern found in this period might stem from a different slope in the increased task-related activation and not from differences in the sustained level of activity (Fig 5). This pattern would align with previous results that support that FPN regions encode evidence accumulation towards a particular decision (Ploran et al., 2011; Gratton et al., 2017). Indeed, in the knapsack task we would expect that lower TCC entails faster evidence accumulation towards a solution.

**Figure 3:**
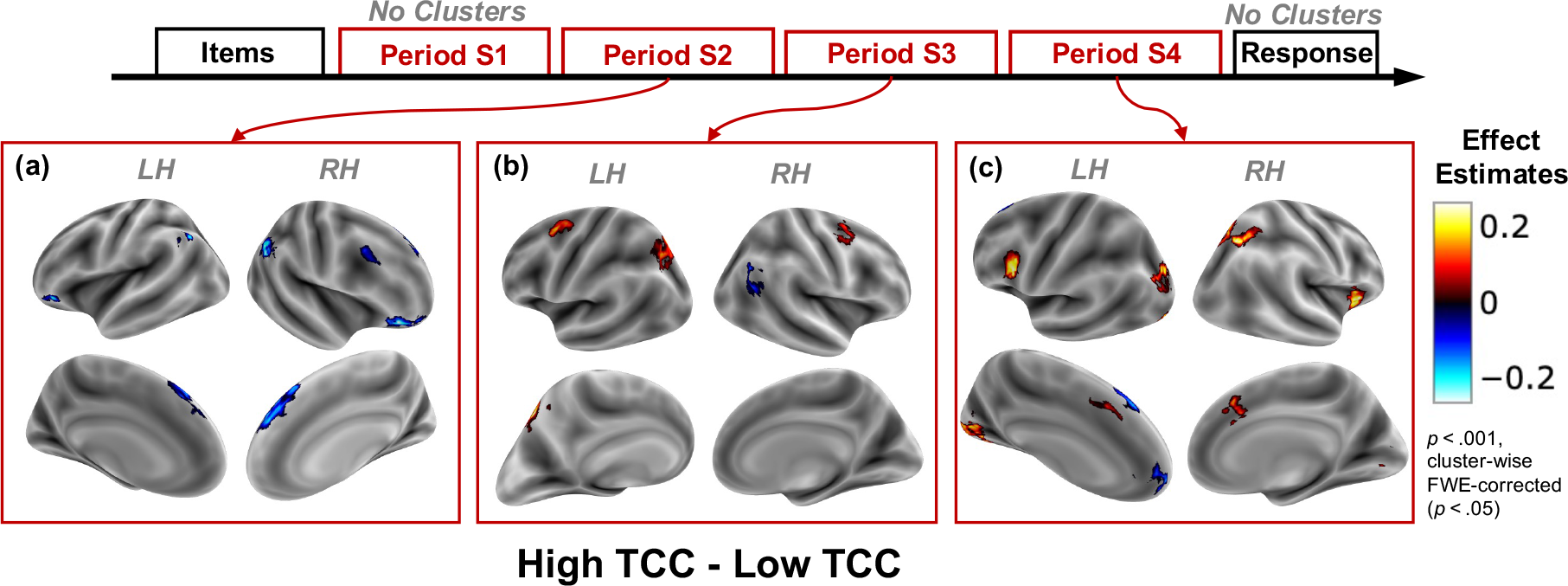
Neural correlates of TCC. Brain activation effect estimates (*β*) for the high vs. low TCC contrast (*β_highT CC_ − β_lowT CC_* ). A positive contrast represents a higher BOLD activity on instances with high TCC compared to low TCC. Significant cluster-wise FWE-corrected (*p <* 0.05) clusters (with an uncorrected threshold of *p <* 0.001) are presented for each of the contrasts estimated using the Boxcar analysis. Each panel represents a different period in the solving stage. **(a)** Period S2, **(b)** period S3, **(c)** period S4. No significant clusters were found for period S1 nor for the response stage parameters.

**Table 1:**
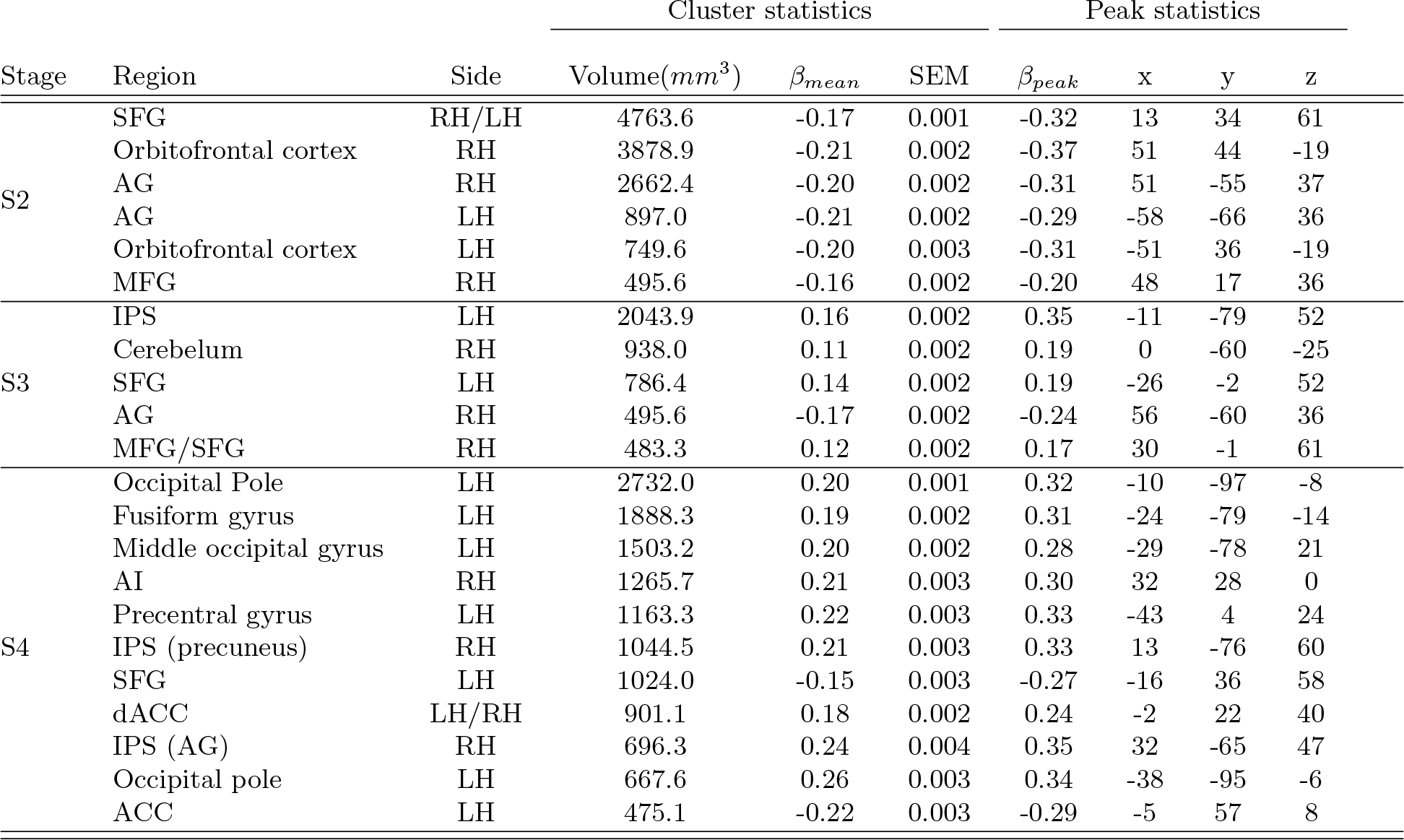
TCC clusters. . Significant cluster-wise FWE-corrected (p < 0.05) clusters (using an uncorrected threshold of p < 0.001) from the High TCC - low TCC contrast. Coordinates are in MNI space.

During the third period of the solving stage, the TCC contrast still showed significant clusters along the FPN, but the pattern overall changed, with respect to period S2. Critically, we found that a different set of regions within the FPN now showed a positive correlation with TCC. Specifically, we found positive clusters in the left SFG, left IPS, the cerebellum as well as a cluster in the right dorsolateral prefrontal cortex (dlPFC) in between the MFG and the SFG. Interestingly, the right AG kept on displaying a negative correlation with TCC during this period.

Finally, during the fourth, and last, period of the solving stage, a new set of clusters was identified. Markedly, this new set of clusters include regions from both CON, FPN as well as significant clusters in the occipital lobe. In general the activation in these clusters activate positively with that of TCC. These include the dACC and right AI from the CON as well as the precentral gyrus and the IPS from the FPN. The right IPS activation is segregated into two clusters, one medial and superior that overlaps with the precuneus and one more lateral that overlaps with the AG. The only two clusters that correlated negatively with TCC are those located in the ACC, as well as a cluster in the left SFG that overlaps with SFG cluster found in the second period.

We did not find any significant clusters during the response stage.

##### Neural correlates of satisfiability

We expected the asymmetry between satisfiable and unsatisfiable instances to reflect differences in control signals associated with reliability. Specifically, we hypothesized that satisfiable instances would be associated with higher reliability, given that once a solution witness is found, verifying that the proposed solution is correct is a polynomial-time operation (tractable problem). In contrast, for unsatisfiable instances, verifying a proof of non-existence is conjectured to be intractable, and thus, computationally harder to verify.

Therefore, we expected regions that have been linked to monitoring of uncertainty to be more active during a trial with an unsatisfiable instance compared to a satisfiable one. In particular, we conjectured higher activation of the CON, on unsatisfiable instances, during late stages of the trial (Neta et al., 2017, 2014; Bossaerts, 2018; Fouragnan et al., 2018).

Interestingly, and contrary to our expectations, we found significant clusters from the first period of the solving stage (Fig 4, Table 2). Moreover, significant clusters did not extend to the response screen, which was also in opposition to our conjecture. Most of the clusters during the solving stage showed a lower BOLD activity for unsatisfiable instances. These clusters extended from period one to period four of the solving stage. Notably, the posterior cingulate showed a lower sustained activation on unsatisfiable instances throughout the solving stage (periods two, three and four). Similarly, different clusters in the SFG had significant clusters throughout the solving stage. Additionally, similar to the clusters found for the TCC contrast, the AG showed bilateral activation during the second period of the solving stage. Interestingly, a bigger AG cluster was found on the left hemisphere compared to the right, in contrast to the right laterality predominance of AG found in the TCC contrast.

**Figure 4:**
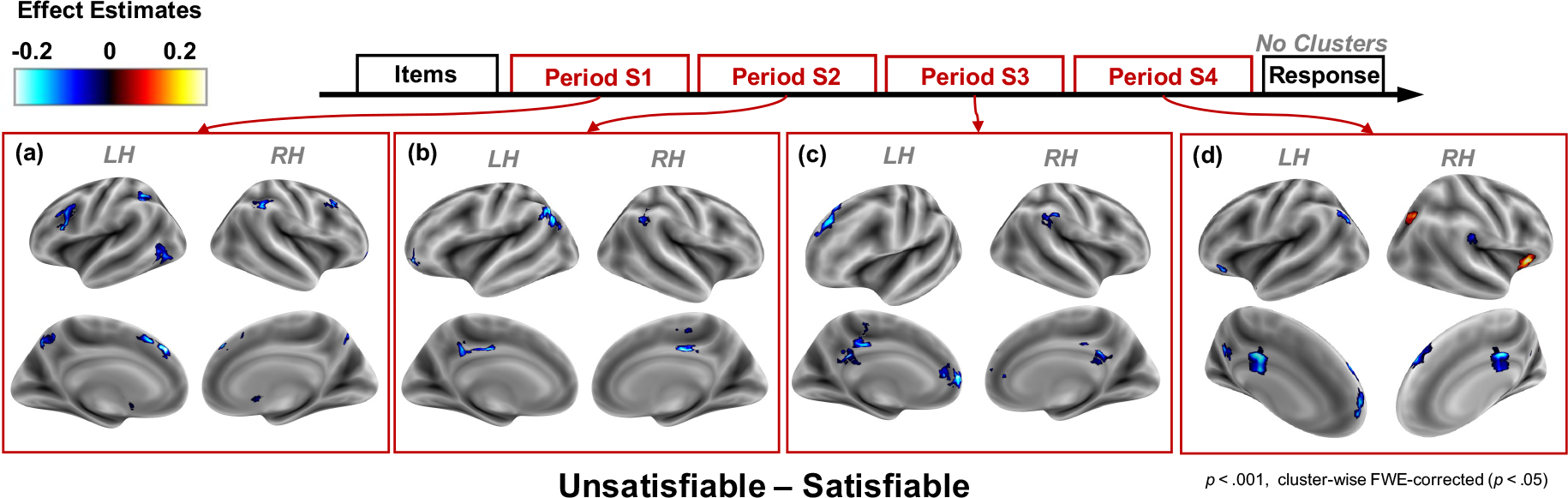
Neural correlates of satisfiability. Brain activation effect estimates (*β*) for the unsatisfiable vs. satisfiable contrast (*β_unsatisfiable_ − β_satisfiable_*). A positive contrast represents a higher BOLD activity on unsatisfiable instances. Significant cluster-wise FWE-corrected (*p <* 0.05) clusters (with an uncorrected threshold of *p <* 0.001) are presented for each of the contrasts estimated using the Boxcar analysis. Each panel represents a different period in the solving stage. **(a)** Period S1, **(b)** period S2, **(c)** period S3 and , **(d)** period S4. No significant clusters were found in the response stage.

**Table 2:** Satisfiability clusters. . Significant cluster-wise FWE-corrected (p < 0.05) clusters (using an uncorrected threshold of p < 0.001) from the Unsatisfiable-Satisfiable contrast. Coordinates are in MNI space.

| Stage | Region | Side | Cluster statistics |  |  | Peak statistics |  |  |  |
| --- | --- | --- | --- | --- | --- | --- | --- | --- | --- |
| | | | Volume($mm^3$) | $\beta_{mean}$ | SEM | $\beta_{peak}$ | x | y | z |
| S1 | SFG | LH | 1876.0 | -0.14 | 0.002 | -0.23 | -3 | 38 | 42 |
|  | Supramarginal gyrus | RH | 1740.8 | -0.13 | 0.002 | -0.25 | 54 | -46 | 56 |
|  | Supramarginal gyrus | LH | 1425.4 | -0.12 | 0.001 | -0.17 | -42 | -47 | 40 |
|  | Inferior occipital cortex | LH | 1159.2 | -0.13 | 0.002 | -0.22 | -61 | -65 | -12 |
|  | MFG | LH | 905.2 | -0.15 | 0.001 | -0.20 | -48 | 18 | 28 |
|  | Caudate | LH | 880.6 | -0.17 | 0.002 | -0.24 | -8 | 2 | -1 |
|  | MFG | RH | 868.4 | -0.12 | 0.003 | -0.23 | 37 | 30 | 53 |
|  | Cerebellum | LH | 667.6 | -0.12 | 0.002 | -0.18 | -38 | -79 | -49 |
|  | Precuneus | LH | 516.1 | -0.19 | 0.003 | -0.26 | -2 | -65 | 44 |
|  | Frontal pole | RH | 450.6 | -0.14 | 0.002 | -0.18 | 29 | 58 | -9 |
|  | Caudate | RH | 450.6 | -0.17 | 0.003 | -0.23 | 8 | 4 | 0 |
| S2 | AG | LH | 2523.1 | -0.19 | 0.002 | -0.29 | -62 | -60 | 29 |
|  | Posterior cingulate | RH | 696.3 | -0.20 | 0.003 | -0.25 | 2 | -25 | 40 |
|  | AG | RH | 585.7 | -0.18 | 0.003 | -0.28 | 62 | -57 | 34 |
|  | SFG | LH | 577.5 | -0.23 | 0.004 | -0.40 | -38 | 62 | -8 |
|  | Posterior cingulate | LH | 479.2 | -0.16 | 0.002 | -0.19 | -10 | -41 | 37 |
| S3 | SFG | LH | 1511.4 | -0.15 | 0.002 | -0.22 | -21 | 36 | 55 |
|  | Anterior cingulate | LH | 708.6 | -0.21 | 0.003 | -0.30 | -5 | 52 | 4 |
|  | Supramarginal gyrus | RH | 696.3 | -0.14 | 0.002 | -0.20 | 64 | -28 | 39 |
|  | Frontal pole | RH | 692.2 | -0.14 | 0.002 | -0.21 | 13 | 58 | 31 |
|  | Anterior cingulate | LH | 593.9 | -0.15 | 0.002 | -0.24 | -6 | 46 | 12 |
|  | Posterior cingulate | LH | 577.5 | -0.17 | 0.002 | -0.22 | -2 | -28 | 45 |
|  | Posterior cingulate | LH | 487.4 | -0.18 | 0.004 | -0.28 | -2 | -44 | 28 |
| S4 | Posterior cingulate | RH | 1384.5 | -0.22 | 0.003 | -0.33 | 0 | -18 | 34 |
|  | SFG | RH | 1306.6 | -0.18 | 0.002 | -0.26 | 14 | 50 | 42 |
|  | AI | RH | 901.1 | 0.25 | 0.003 | 0.36 | 32 | 28 | 0 |
|  | AG | LH | 659.5 | -0.24 | 0.003 | -0.30 | -48 | -68 | 44 |
|  | SFG | LH | 647.2 | -0.17 | 0.004 | -0.26 | -14 | 52 | 40 |
|  | Precuneus | LH | 581.6 | -0.19 | 0.005 | -0.27 | -5 | -57 | 31 |
|  | Occipital superior cortex | RH | 544.8 | 0.17 | 0.005 | 0.27 | 29 | -63 | 36 |
|  | SFG / Frontal pole | LH | 512.0 | -0.24 | 0.003 | -0.34 | -3 | 65 | 16 |
|  | Orbitofrontal cortex | LH | 454.7 | -0.23 | 0.003 | -0.33 | -46 | 28 | -20 |
|  | Supramarginal | RH | 438.3 | -0.12 | 0.002 | -0.18 | 54 | -33 | 32 |

The only two clusters that showed a significantly higher activity on unsatisfiable instances were the right AI and the occipital superior cortex, both present only during period four of the solving stage. The significant cluster found in the AI is in line with our hypothesis that unsatisfiable instances are related to higher markers of uncertainty, a signal which we expected to find in the CON. However, in disagreement with our hypothesis, we did not find a significant satisfiability cluster in the dACC. This may be due to insufficient statistical power of the whole brain analysis (see Fig 5).

**Figure 5:**
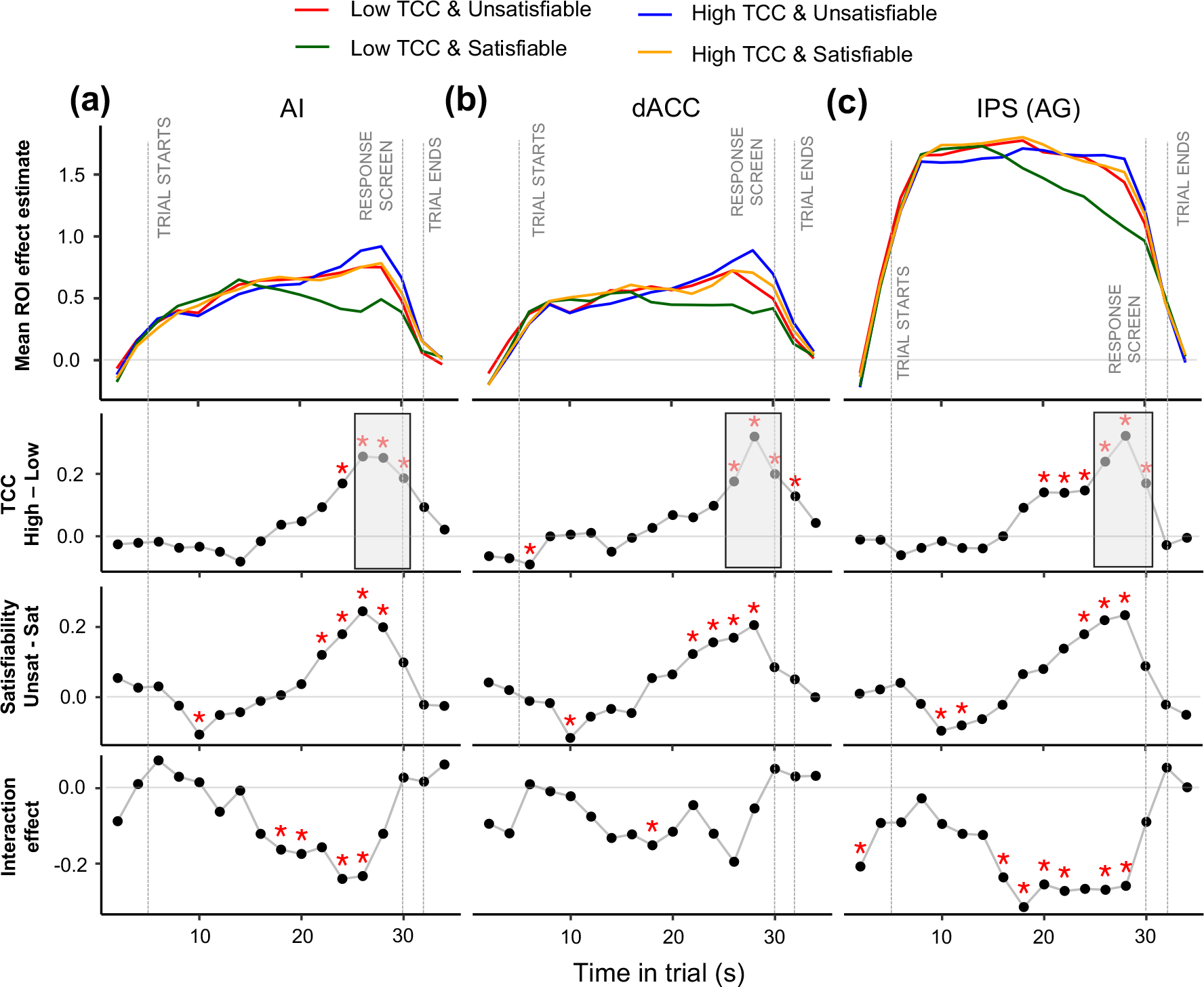
Temporal dynamics of BOLD in regions of interest. Mean effect estimate (*β*) for each ROI against time in trial. The effect at each time point represents the mean *β_F IR_* over all of the voxels from each ROI: right AI **(a)**, dACC **(b)**, and right IPS cluster extending to the angular gyrus **(c)**. In the top row of figures the *β_F IR_* ’s characterize the coefficients of an FIR regression with four conditions: satisfiability*×*TCC. The *β_F IR_* parameters are aligned to the BOLD signal, which has a lag with respect to the task time. To correct for this, the gray time-markers represent the task-events by assuming a 5 seconds BOLD signal lag. In the second row, the TCC contrast (*β_high_ − β_low_* ) is presented. The third row displays the satisfiability contrast (*βunsat − βsat*). The bottom row shows the interaction effect between TCC and satisfiability ([*β_highT CC,unsat_ − β_lowT CC,unsat_*] *−* [*β_highT CC,sat_ − β_lowT CC,sat_*]). Red asterisks represent significance at a 0.05 significance level. Significance levels in the gray shaded regions are suggestive only; they represent the time period and contrast from which the ROIs were selected.

##### Neural correlates of accuracy

Although participants did not receive any feedback during the task, we expected to see error-related signals late in the trial. Although these signals would not represent the integration of novel exogenous information (since there was no feedback), we conjectured that participants would hold a subjective belief of the expected accuracy (or reward) of their answer (e.g, Duverne and Koechlin (2017)). In line with our hypothesis, we found that activity in both FPN and CON was positively correlated with erring during the response stage (Appendix E).

##### ROI dynamics

Three ROIs were selected (see Section 4.9.3) to investigate more closely the neural dynamics of computational complexity. We included in our analysis the dACC due to its proposed involvement in the allocation of control (Shenhav et al., 2013; Dosenbach et al., 2006; Silvetti et al., 2018; Vassena et al., 2017; Holroyd and Yeung, 2012; Alexander and Brown, 2011) as well as the right AI because of its involvement in encoding control signals and uncertainty in particular (Neta et al., 2017, 2014; Bossaerts, 2018; Fouragnan et al., 2018). In order to compare the neural activity in these regions, which are generally attributed to control, with relevant processing units, we selected a region associated with mathematical calculations, the right IPS (Matejko and Ansari, 2018; Brannon, 2006; Arsalidou and Taylor, 2011).

We explored the BOLD effect estimates (*β_F IR_*) for the 2*×*2 balanced factorial design (satisfiability*×*TCC) employing Finite Impulse Response (FIR) analysis at one-second resolution (see Section 4.9.4). We found similar patterns in AI and dACC. In both regions, the BOLD signal rose throughout the task and quickly decreased around the time the solving stage ended (Fig 5). The activity pattern in the IPS showed a different pattern to that of CON regions. In this region, the BOLD signal increased quickly early on in the trial and was sustained until it started decreasing later on in the trial. The moment at which the decrease started was modulated by TCC and satisfiability (Fig 5c).

Interestingly, there seemed to be an interaction between satisfiability and TCC (Fig 5 fourth row of panels). Specifically, satisfiable instances with low TCC started showing a decrease in activity from early on in the trial on all three regions (Fig 5 green line). Conversely, unsatisfiable instances with high TCC showed a more pronounced peak late in the trial in both AI and dACC (blue line).

When contrasting the effect of TCC in each of the ROIs, we find that there is a significant positive effect of TCC from mid-way through the trial in the right IPS/AG (Fig 5 second row of panels). This differs from the results obtained in the whole brain analysis. Similarly, when estimating the effect of satisfiability (Fig 5 third row of panels), the results marginally differ from those of the whole-brain analysis. Firstly, the ROI analysis reveals that there is an effect of satisfiability on all three regions late in the solving-stage. Secondly, the effect of satisfiability starts in the AI and dACC mid-way through the trial. Interestingly, the effect of TCC seems to precede that of satisfiability in the IPS, whereas in the dACC the effect of satisfiability seems to precede that of TCC.

Altogether, these results suggest that both satisfiability and TCC correlate with activity in all three regions, but that their effect might have different neural temporal signatures. Importantly, the sign of the effect was in line with our hypothesis: a higher signal in these regions was generally related to higher TCC and unsatisfiability. The only exceptions happen briefly early on in a trial.

Finally, we explored the interaction between neural markers of accuracy and the proposed metrics of computational difficulty (Appendix E). To do this, we analyzed (employing FIR in the three ROIs) the neural markers of accuracy separately for each type of instance (i.e., level of TCC and satisfiability). Overall, our results suggest a link between neural markers of accuracy and metrics of computational difficulty. Specifically, we found a significantly lower BOLD signal on incorrect instances with low TCC and on incorrect satisfiable instances. This effect was particularly evident in the IPS and marginally in the AI. Importantly, these trends showed from the moment the trial started, suggesting that for instances associated with low computational difficulty (i.e., low proof hardness and low complexity), the accuracy could be predicted from early on in the trial from BOLD activity in the IPS.

##### Psychophysiological interactions (PPI)

To study the effect of TCC and satisfiability on *functional connectivity* we conducted a PPI analysis to gauge the functional synchronization between each of the ROIs and other regions in the brain. Explicitly, we performed whole-brain PPI analyses employing the three considered ROIs (dACC, rAG and rAI) as seed regions. For these regressions we modeled the task (items and solving stages together) with two boxcar functions of equal length (12.5s) (Fig 6). This allowed us to study PPI task interactions separately for an early period (PPI-1: first 12.5 seconds of the task) and a late period (PPI-2: last 12.5 seconds). We found a similar and generalized pattern of connectivity for all three ROIs and both periods when contrasting the PPI effect compared to baseline (Appendix Fig 8). This suggest that the task has a similar effect on the BOLD synchronization between the three ROIs and several regions.

**Figure 6:**
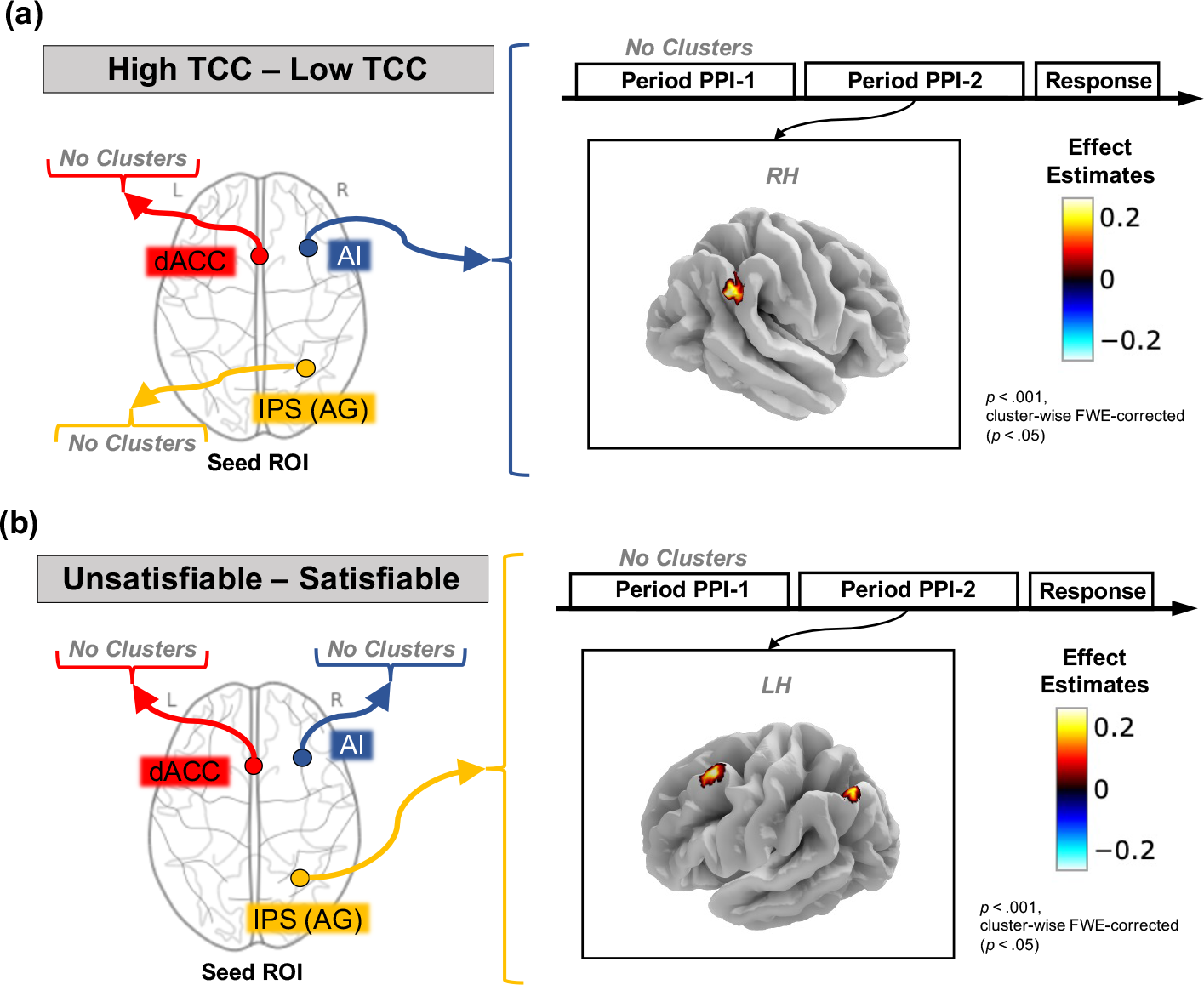
PPI results. The effect of instances’ properties on connectivity: **(a) TCC**, **(b) Satisfiability**. The left panel represents the seed region used for the analysis (dACC, rAG or rAI). The right panel shows the clusters that display a significant PPI connectivity effect for a particular seed region and period. *Significant cluster-wise FWE-corrected (p <* 0.05*) clusters (with an uncorrected threshold of p <* 0.001*) are presented. “No clusters”: No significant clusters were found for this analysis*.

When comparing the connectivity between instances with high and low TCC, we found one significant cluster with differential connectivity. This cluster, located along the rAG and the supramarginal gyrus, showed a change in connectivity to the rAI (seed region) between high and low TCC instances during the second PPI period (Fig 6a; Table 3). We also explored the differences in the PPI connectivity between unsatisfiable and satisfiable instances. We observed a significant PPI effect of satisfiability between the right IPS/AG (seed) and the left MFG, as well as with the left AG, during the second PPI period (Fig 6b; Table 3). Overall, these results suggest that instance properties have an effect on the synchronicity between the ROIs and a limited collection of clusters. However, this effect is only significant during the later part of the solving stage.

**Table 3:** PPI clusters. . The effect of instances’ properties on connectivity. Significant cluster-wise FWE-corrected (p < 0.05) clusters (using an uncorrected threshold of p < 0.001). Coordinates in MNI space.

| Contrast | Region | Side | Cluster statistics |  |  | Peak statistics |  |  |  |
| --- | --- | --- | --- | --- | --- | --- | --- | --- | --- |
| | | | Volume($mm^3$) | $\beta_{mean}$ | SEM | $\beta_{peak}$ | x | y | z |
| TCC | AG/Supramarginal Gyrus | RH | 573.4 | 0.26 | 0.004 | 0.36 | 61 | -50 | 36 |
| Satisfiability | MFG | LH | 499.7 | 0.300 | 0.007 | 0.475 | -43 | 18 | 59 |
|  | AG | LH | 483.3 | 0.266 | 0.006 | 0.432 | -54 | -65 | 47 |

##### Granger causality analysis

In order to explore *effective connectivity* patterns, we studied temporal directionality with Granger Causality (GC). We were particularly interested in the effect of complexity and proof hardness on the effective connectivity between our three ROIs. We found a significant bidirectional connectivity between all the ROIs at baseline during the experiment. Additionally, during the solving stage, we found that there was a significant change in GC from dACC to rAG. However, we did not find any significant changes in the effective connectivity between high and low TCC instances nor between unsatisfiable and satisfiable instances. Taken together, these results suggest that the effects of TCC and satisfiability on neural activity propagate through the ROIs via baseline effective connectivity (present during the solving stage) and not through a direct effect on the effective connectivity (see Appendix E.2).

## 3. Discussion

The study of the neural underpinnings of problem-solving has, to date, been centered on tractable problems. This line of research has led to the characterization of networks and processes that support problem-solving. A critical shortcoming of existing studies is the lack of a formal and generic characterization of cognitive resource requirements, which is particularly problematic in relation to intractable problems where problems require an extended period of time to solve and involve an extensive search space (e.g., MacGregor and Chu, 2011; Acuña and Parada, 2010; Murawski and Bossaerts, 2016; Franco et al., 2021b). Here, we present a framework, grounded in computational complexity theory, to study the neural underpinnings of problem-solving that overcomes previous limitations.

We empirically test this framework in the knapsack decision task using ultra-high field fMRI. Our findings shed light into the neural processes supporting problem-solving. Firstly, our findings not only extend but solidify the research on the neural correlates of cognitive demand by exploring the processes associated with one specific dimension of cognitive demand: computational complexity. Importantly, this is done in a task-independent way in the sense that these metrics can be applied to a whole class of problems (i.e., NP-complete). Secondly, rather than studying cognitive demand starting from “what people are thinking,” we rely on the theory of computational complexity to identify intrinsic properties of a problem to delineate cognitive requirements. These intrinsic properties allowed us to discover relevant neural markers and their dynamics, similar to how risk and variance have been shown to affect decisions in probabilistic tasks (D’Acremont and Bossaerts, 2008). Finally, the results presented here complement the investigation of cognitive control by providing a framework that can be employed to extend previous findings to tasks that involve intractable problems. Critically, cognitive control involves the dynamic allocation of cognitive resources that stem from an interaction between the cognitive requirements of a task and the resources available. The framework put forward here provides a theoretical foundation for the characterization of the former.

Extensive research has studied the neural correlates of cognitive demand. This program has characterized the MDS, a network of regions that respond robustly to cognitive demand irregardless of the task at hand (Fedorenko et al., 2013; Assem et al., 2020; Duncan and Owen, 2000; Crittenden et al., 2016). This has been done using several tasks including perceptual target detection and memory retrieval, among many others. Notably, most of the tasks employed to date have been based on tractable problems. Moreover, many of the tasks employed modulate cognitive demand of the task by tuning the amount of processing needed on one specific dimension of cognitive processing. For instance, in perceptual tasks, signal to noise ratio is modulated (e.g., Aben et al., 2020; Dubis et al., 2016; Hanks and Summerfield, 2017; Ploran et al., 2011); alternatively, in memory retrieval tasks, the amount of information to be stored/retrieved is tuned (e.g., Gratton et al., 2018; Fedorenko et al., 2013). The lack of a generic (problem-independent) definition of cognitive demand hinders the generalization of this approach to new problems. Importantly, the level of cognitive demand might be highly related to the strategies used. Here, we propose a way forward, grounded in the assumption that hardness is, at least partially, an intrinsic characteristic of the problem at hand.

Following this approach, we operationalized cognitive demand via TCC and found that the neural correlates of TCC overlapped with those associated with the MDS. In particular, the positively correlated clusters (higher activation in high TCC instances) in the FPN and CON resembled those of the MDS. Notably, we found clusters in the AI, the dACC, the precentral gyrus and the IPS, which have been associated with the MDS (Fedorenko et al., 2013). Importantly, our results display a dynamic process in which the neural correlates of TCC vary throughout the different stages of the task. This suggests that the MDS can be construed as a heterogeneous set of regions that play a dynamic and varying role at different stages in problem-solving.

It is worth highlighting that we are not arguing for the proposed framework to replace other methodological approaches in the study of cognitive demand. Instead, we assert that both approaches complement each other. Critically, complex tasks involve the interplay of several computational processing units such as working memory, logical operations, processing of numerical magnitudes among many others. Our approach, as it stands, is not able to differentiate among these sub-processes. A proper understanding of problem-solving requires both the study of these sub-processes independently, like in more classical approaches (e.g., Gratton et al., 2018), as well as in tandem in order to understand how they interact to support computationally hard problem-solving, like done in this paper.

A related effect of these properties on neural processes is through the encoding of relevant task markers that could be employed during problem-solving (Yoo et al., 2021; Koechlin, 2016). These neural markers include markers of performance such as expected error (Neta et al., 2017; Bossaerts, 2018), variance in this expectation (uncertainty) (Neta et al., 2017, 2014; Bossaerts, 2018), as well as markers that encode the evidence towards a particular response (Ploran et al., 2011; Gratton et al., 2017) or even the merit of alternative strategies (Duverne and Koechlin, 2017; Donoso et al., 2014). Critically, we made three conjectures with regards to these neural markers. First, we hypothesized to see markers of performance, related to TCC, from early on in the trial. Second, we conjectured we would see markers of reliability, related to satisfiability, in regions shown to encode uncertainty. Third, we expected to find neural correlates of accuracy late in the trial, which would be associated to expected performance.

With regards to our first hypothesis, we argue that TCC is a feasible metric that can be related to markers of performance and efficacy of effort from early on in the trial. Firstly, TCC has been shown to be correlated with human performance (Franco et al., 2021b). Secondly, TCC can be potentially estimated from early on in the solving stage without the need to know the solution to the problem. As such, we expected to see neural correlates of TCC from early on in the trial. Specifically, we expected to see markers of TCC from early on in the solving stage in the CON (Shenhav et al., 2013; Bossaerts, 2018; Neta et al., 2014). Contrary to our expectations, we only found significant clusters in the CON starting from the third period of the solving stage. These might reflect markers of expected performance, but other explanations cannot be excluded. For instance, this effect might reflect differences in time-on-task between TCC conditions (Grinband et al., 2011). This explanation, however, would still allow these activation patterns to represent differences in neural markers such as reliability and expected performance. This follows from the fact that time-on-task is an endogenous variable of the system. That is, the agent decides when to stop reasoning about the problem, and as such, this decision would likely follow from a subjective belief on how well they can expect to perform given the current candidate solution. Therefore, differences in time-on-task between high and low TCC instances may be related to differences in subjective beliefs of both expected performance and reliability.

Additional to the reported clusters that correlated positively with computational complexity, we found a set of clusters that correlated negatively with TCC. These clusters are concentrated in the second period of the solving stage, but are also found in the third and fourth periods of the solving stage. These results might be explained by the encoding of evidence accumulation signals (Ploran et al., 2011). Arguably, evidence toward a solution can be accumulated faster in low TCC compared to high TCC instances. This would imply that regions that encode evidence accumulation would show a higher activation on low TCC instances early in the trial, in accordance with the pattern found on the second period of the solving stage.

Turning now to our second conjecture regarding the neural markers of reliability, we explored the correlates of satisfiability during problem-solving. We expected to see activation related to satisfiability in regions previously associated with uncertainty encoding, specifically, in the CON. In line with our hypothesis, we found a significant positive relation between unsatisfiability and activity in the CON that started halfway through the solving stage. In line with our conjecture, we found that the neural markers of satisfiability overlap with regions that encode probabilistic uncertainty. This suggests that reliability and uncertainty might constitute analogous constructs that are encoded similarly across tasks and that could serve a generic role in decision-making.

Contrary to our expectations, we found several regions that displayed an increase in activity during satisfiable instances from early on in the solving stage. This result is perplexing because knowing the satisfiability of the problem equates to having solved the problem, which would not be expected early on in the trial. A possible explanation for this is that the clusters found encode evidence accumulation (Ploran et al., 2011; Gratton et al., 2017) and that accumulating evidence towards the solution in satisfiable instances occurs at a different rate than in unsatisfiable instances. Relatedly, these activation patterns might reflect the use of different strategies. However, this account would still require participants to be implementing different strategies, based on satisfiability, as early as during the first few seconds of the solving stage. Overall, future research should attempt to disentangle the effect of proof hardness and the satisfiability of instances. This could be done, for instance, through experiments aimed at testing more nuanced metrics of proof hardness.

Moving on now to consider our conjecture related to the neural markers associated with erring, we explored the effect of accuracy on the neural activation throughout the task. It has been proposed that FPN and CON regions encode task signals related to error detection and error expectation (Neta et al., 2014, 2017; Dosenbach et al., 2006). We hypothesized that participants would represent a subjective belief on the expected accuracy (or reward) of their answer (e.g, Duverne and Koechlin (2017)). In line with our conjecture, we found that activity in both the FPN and CON was positively correlated with erring during the response stage (Appendix E).

Overall, we found evidence that suggests the existence of neural markers related to computational complexity, proof hardness and performance. Taken together, the framework put forward here provides a way to study neural markers associated to subjective beliefs during problem-solving. It is worth noting, however, that while we modulated complexity and proof hardness, many other complexity-related features might be relevant, including, for example, the size of the problem at hand (e.g., Carruthers et al., 2012; MacGregor and Chu, 2011; Dry et al., 2006; van Opheusden and Ma, 2019; Stazyk et al., 1982; De Visscher and Nöel, 2014). Further work in this area is needed to understand the interaction between different sources of computational difficulty in human problem-solving.

Finally, to explore the dynamics related to control during complex problem-solving, we analyzed the functional interaction during problem-solving of three ROIs, two of which have been associated with cognitive control (i.e., CON) and one region which has been associated with processes that were deemed highly relevant for the task at hand (i.e., IPS). We studied synchronization of signals (employing PPI analysis) and explored their effective connectivity (using GC analysis).

Our results support the view that there is a generalized change in signal synchronization during the solving stage compared to baseline. Moreover, when exploring the link between instance properties and synchronicity between regions, we found several clusters whose connectivity was modulated by either satisfiability or TCC. These effects were only present late in the trial. Specifically, we found that TCC modulated the synchronicity between the rAI and the rIPS. Additionally, satisfiability modulated the functional connectivity between the right IPS and two clusters in the left hemisphere, one in the AG and one in the MFG. Overall, these results suggest a differential recruitment of regions during the task, partially modulated by task properties late in the trial. Interestingly, the significant clusters identified in this analysis have been implicated in the performance of mathematical calculations (Arsalidou and Taylor, 2011; Grabner et al., 2009), suggesting that they could support moment-to-moment implementation of strategies. Further work would be needed in order to asses whether the relation, found here, between instance properties and functional synchronization is associated to the implementation of different strategies.

Additionally, we found that the effective connectivity pattern was impervious to the level of TCC and satisfiability. This suggests that the effect of computational complexity on control would occur by generating differential levels of activity within the regions of interest and not via modulation of the effective connectivity between these regions. It is worth noting that the failure to reject the null hypothesis (of no effects of TCC and satisfiability on effective connectivity), however, could be due to lack of power or due to the exclusion of relevant ROIs from the analysis. Further research is needed to assess whether whole brain effective connectivity patterns are affected by computational complexity.

Humans are constantly solving problems that vary in their complexity, ranging from perceptual tasks, such as motion detection and face recognition, to reasoning tasks such as choosing an investment portfolio. Understanding how complexity of these problems affects the neural processes involved in problem-solving is of crucial importance for the understanding of human decision-making. Here, we present a framework that allows for the study of computational difficulty of human problem-solving. We applied this framework and identified a dynamic set of regions in which activation was modulated by different properties related to computational complexity. Overall, our findings provide support to the premise that computational complexity theory, as applied here, provides a useful characterization of cognitive demand and reliability for the study of problem-solving in neuroscience.

## Acknowledgments

This research is supported by a University of Melbourne Graduate Research Scholarship from the Faculty of Business and Economics. Bossaerts acknowledges financial support through a R@MAP Chair from the University of Melbourne.

## Materials and methods

### 4.1. Ethics statement

The experimental protocol was approved by the University of Melbourne Human Research Ethics Committee (Ethics ID 1749616.3). Written informed consent was obtained from all participants prior to commencement of the experimental sessions. Experiments were performed in accordance with all relevant guidelines and regulations.

### 4.2. Participants

Twenty right-handed volunteers from Melbourne University and the surrounding community took part in the study (14 female, 5 male, 1 other; age range = 18-35 years, mean age = 26.6 years). Inclusion was based on age (minimum = 18 years, maximum = 40 years) and on right-handedness. Each participant performed the knapsack decision task in the scanner and performed outside the scanner the knapsack optimization task and a set of basic cognitive function tasks.

### 4.3. Knapsack decision task

In this task, participants were asked to solve a number of instances of the (0-1) knapsack decision problem (Fig 1). In each trial, they were shown a set of items with different values and weights as well as a capacity constraint and a target profit. Participants had to decide whether there exists a subset of those items for which (1) the sum of weights is lower or equal to the capacity constraint and (2) the sum of values yields at least the target profit.

Each trial had four stages. In the first stage (items stage; 3 seconds), only the items were presented. Item values, in dollars, were displayed using dollar bills and weights, in grams, were shown inside a black weight symbol. The larger the value of an item, the larger the dollar bill was in size. Similarly, the larger the weight of an item, the larger its weight symbol was in size. At the center of the screen, a green circle indicated the time remaining in this stage. In the second stage (solving stage; 22 seconds), target profit and capacity constraint were added to the screen inside the green timer circle. In the third stage (response stage; 2 seconds), participants saw a ‘YES’ and a ‘NO’ button on the screen, in addition to the timer circle, and made a response using the keyboard (Fig 1). Finally, a jittered inter-trial rest period of 8, 10 or 12 seconds was shown before the start of the next trial.

Participants completed 56 trials (7 blocks of 8 trials), each showing a different instance of the knapsack decision problem. The order of instances was randomized across participants. The side of the ‘YES’ and ‘NO’ buttons was also randomized.

### 4.4. Instance sampling

Instances were sampled following a 2*×*2 balanced factorial design for the factors TCC (high and low) and satisfiability (satisfiable and unsatisfiable). Specifically, instances selected were sub-sampled from those employed in a previous behavioral study (Franco et al., 2021b). Instances in their study were selected such that *α_c_* was fixed (*α_c_ ∈* [0.40, 0.45]) and the instance constrainedness varied according to *α_p_*. 18 satisfiable instances were selected from the under-constrained region (*α_p_ ∈* [0.35, 0.4]; *low TCC* ) and 18 unsatisfiable instances from the over-constrained region (*α_p_ ∈* [0.85, 0.9]; *low TCC* ). Additionally, 18 satisfiable instances and 18 unsatisfiable instances were sampled near the satisfiability threshold (*α_p_ ∈* [0.6, 0.65]; *high TCC* ). Half of the instances with high TCC were forced to have high/low computational requirements (top/bottom 50%), according to an algorithm-specific ex-post complexity measure of a widely-used algorithm (Gecode ; Gecode Team (2006)). All instances in the experiment had *N* = 6 items and *w_i_*, *v_i_*, *c* and *p* were integers. In the current study we randomly selected 56 of the 72 instances sampled in Franco et al. (2021b). Sub- sampling without replacement was done ensuring that the same number of instances were selected across TCC and satisfiability conditions. Moreover, instances with high TCC were balanced to require high/low computational requirements according to the same algorithm-specific complexity measure employed in their study (i.e., Gecode propagations).

### 4.5. Complementary tasks

Participants were presented a set of complementary tasks outside of the scanner. They were asked to solve a number of instances of the (0-1) knapsack optimization problem. Similar to the knapsack decision task, participants were shown a set of items with different weights and values as well as a capacity constraint. However, unlike the decision variant, no target profit was presented. Participants had to find the subset of items that *maximized* total value subject to the capacity constraint (see Appendix C.2).

We also tested participants’ performance on five aspects of cognitive function that we considered relevant for the knapsack tasks, namely, working memory, episodic memory, strategy use, processing and psychomotor speed, as well as mental arithmetic. To do so, we administered a set of tasks from the Cambridge Neuropsychological Test Automated Battery (CANTAB; Appendix D). Specifically, we asked participants to perform the Reaction Time (RTI), Paired Associates Learning (PAL), Spatial Working Memory (SWM) and Spatial Span (SSP). In addition, participants were presented with a set of mental arithmetic problems (Appendix D).

### 4.6. Procedure

Participants were asked to fill in an MRI screening form before attending the experiment. Once at the experiment, participants were presented with a plain language statement and a consent form. After reading these and providing written informed consent, participants were instructed in the tasks and completed a practice session of the knapsack decision task. Participants then underwent an MRI safety check and debriefing.

Before being scanned, participants solved the CANTAB RTI task outside of the scanner. This was followed by the scan session in which they performed the knapsack decision task. Afterwards, outside of the scanner, they completed the CANTAB RTI task again, followed by the knapsack optimization task. Subsequently, they completed the remaining CANTAB tasks in the following order: PAL, SWM and SSP. Finally, they performed the mental arithmetic task and completed a set of demographic and debriefing questionnaires. Altogether, the experimental session lasted around three hours.

Participants received a show-up fee of A$10, as well as monetary compensation based on performance. They earned A$1.2 for each correct answer in the knapsack decision task and for each correct answer in the knapsack optimization task.

### 4.7. Behavioral statistical analyses

The R programming language was used to analyze the behavioral data. All of the linear mixed models (LMM), generalized logistic mixed models (GLMM) and censored linear mixed models (CLMM) included random effects on the intercept for participants (unless otherwise stated). Different models were selected according to the data structure. GLMM were used for models with binary dependent variables, LMM were used for continuous dependent variables and CLMM were used for censored continuous dependent variables (e.g., time-on-task).

All of the models were fitted using a Bayesian framework implemented using the probabilistic programming language Stan via the R package ‘brms’ (Bürkner, 2017). Default priors were used. All population-level effects of interest had uninformative priors; i.e., an improper flat prior over the reals. Intercepts had a student-t prior with 3 degrees of freedom and a scale parameter that depended on the standard deviation of the dependent variable after applying the link function. The t-student distribution was centered around the mean of the dependent variable. Sigma values, in the case of Gaussian-link models, had a half student-t prior (restricted to positive values) with 3 degrees of freedom and a scale parameter that depended on the standard deviation of the dependent variable after applying the link function. Standard deviations of the participant-level intercept had a half student-t prior that was scaled in the same way as the sigma priors.

Each of the models presented was estimated using four Markov chains. The number of iterations per chain was by default set to 2000. This parameter was adjusted to 4000 on some models to ensure convergence, which was verified using the convergence diagnostic *R*^^^. All models presented reach an *R*^^^ *≈* 1.

Statistical tests were performed based on the 95% credible interval estimated using the highest density interval (HDI) of the posterior distributions calculated via the R package ‘parameters’ (Lüdecke et al., 2020). For each statistical test we report both the median (*β*_0.5_) of the posterior distribution and its corresponding credible interval (*HDI*_0.95_).

No participant nor trial was excluded from the data analysis of the knapsack decision task.

### 4.8. MRI data acquisition

We collected the fMRI images using a 7 Tesla Siemens MAGNETOM scanner located at the Melbourne Brain Centre (Parkville, Victoria) with a 32-channel radio frequency coil.

The BOLD signal was measured using a multiband echo-planar imaging sequence (TR = 800 ms, TE = 22.2 ms, FA = 45°). We acquired 84 interleaved slices (thickness = 1.6 mm, gap = 0 mm, FOV = 208 mm, matrix = 130x130, multi-band factor = 6, voxel size=1.6×1.6×1.6mm^3^) per volume. 380 volumes were acquired on each run while recording cardiac and respiratory traces.

After five functional runs (one resting state run followed by four task runs), a high resolution (0.7 mm isotropic) anatomical image was acquired using an MP2RAGE pulse sequence (TR=5000 ms, TE=3.07 ms, TI1 = 700ms, FA1 = 4°, TI2 = 2700ms, FA1 = 5°, matrix=330×330, voxel size=0.73×0.73×0.73mm^3^,

FOV=240 mm, 224 slices, slice thickness = 0.73). Afterwards, another three functional runs were performed, followed by a diffusion weighted imaging (DWI) multi-band sequence (TR=7000 ms, TE=72.4 ms, FA =90°, FoV = 210 mm, matrix = 170x170, slice thickness =1.24, voxel size = 1.24*m*^3^, 128 slices, multi-band factor=2).

### 4.9. Imaging statistical analyses

#### 4.9.1. Preprocessing

Initial preprocessing of the data was performed using AFNI (Cox, 1996) and the Advanced Normalization Tools (ANTs) software. For each subject, pulse and cardiac noise was regressed out from the functional scans. These were then slice-time corrected and the volumes were motion-corrected by registering them to the first volume of the first functional run. The mean image of the first run was co-registered to the anatomical scan (down-sampled) and this transformation was applied to all of the functional volumes. Afterwards, each participant’s anatomical scan was used for calculation of transformation parameters to normalize the functional images into the Montreal Neurological Institute (MNI) space (see Appendix A for more details).

#### 4.9.2. Whole-brain analysis (boxcar)

Whole-brain analyses were performed by fitting generalized linear models (GLM) using AFNI (Cox, 1996). Before the regressions were implemented, we spatially smoothed the functional volumes with a 4.8mm FWHM Gaussian kernel. Additionally, volumes with motion or signal outliers were censored from each of the regressions.

We performed GLM regressions to explore three contrasts of interest. Specifically, we tested the neural correlates of TCC (high TCC vs. low TCC), satisfiability (unsatisfiable vs. satisfiable) and accuracy (correct vs. incorrect). In each of the regressions the solving phase (22s) was modeled using four boxcar functions of equal duration (5.5s):

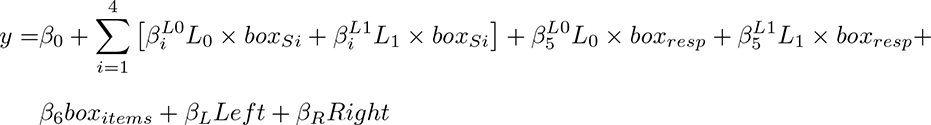

where *L*_0_ and *L*1 correspond to the different levels of interest (e.g., high TCC and low TCC respectively) and *box_Si_*, *box_resp_* and *box_items_* correspond to the boxcar functions of the solving, response and items stages, respectively. *Left* and *Right* correspond to the button pressed by the participant.

Group level analyses were performed using mixed effects multilevel modeling (Chen et al., 2012). All whole-brain analysis results are reported with a clusterwise threshold of *p <* 0.05 corrected for multiple comparisons across the whole brain, using an uncorrected voxelwise threshold of *p <* 0.001.

#### 4.9.3. ROI specification

We were particularly interested in how control and subjective beliefs of cognitive demand and reliability were involved in complex problem-solving. To study these dynamic processes we selected three regions of interest (ROIs) that have been implicated in the processes of interest. Firstly, we included in our analysis the CON (dACC and AI) due to its proposed involvement in the allocation of control (Shenhav et al., 2013; Dosenbach et al., 2006; Silvetti et al., 2018; Vassena et al., 2017; Holroyd and Yeung, 2012; Alexander and Brown, 2011) and uncertainty encoding (Neta et al., 2017, 2014; Bossaerts, 2018), which we conjectured would be highly related to encoding of reliability. Secondly, we included a region that has been involved in moment-to-moment processing operations during problem-solving. We expected the knapsack task to engage processing units associated with number processing and mathematical calculations. Therefore, we selected a region that has been widely connected to ‘processing’ in mathematical problem-solving, the right IPS (Matejko and Ansari, 2018; Brannon, 2006; Arsalidou and Taylor, 2011).

The three ROIs were selected from the clusters found when contrasting high and low TCC in the last boxcar during the solving stage (period S4). We chose the contrast for the fourth boxcar for a few reasons. We expected that during this last period of the solving stage we would be able to see a marked differentiation in the cognitive demand between instances with high and low TCC. We expected instances with low TCC to require less computational time and thus, we hypothesized that, on average, participants would be still making calculations during the period S4 for high TCC instance, but not for low TCC instances. This was further indicated by a parallel pilot study that found that participants spent on average 17.9s solving an instance with low TCC and 21.2s on those with high TCC (period S3 ends at 19.5s of solving stage). Importantly, we believed that these differences in cognitive demand would be reflected as well in a differentiation in the control activity in the system. Critically, we expected the monitoring of control variables such as expected performance would differ between types of instances. For instance, we expected the subjective markers of performance would converge to actual performance levels in the late stages of the solving stage (Franco et al. (2021b); Fig 2), which would imply higher subjective beliefs of expected performance for low TCC. Additionally, we expected that this contrast would allow us to control for task-set signals (Dosenbach et al., 2007). We conjectured that the task-set signals would be maintained during the whole solution-stage, so the proposed contrast would not capture task-set signals encoding goals nor the underlying structure of the task.

Among the significant clusters found around the right IPS, we chose the IPS (AG) cluster (peak: x=32, y =-65, z=47) because of its overlap with the regions that were found to be associated with mathematical calculations in the meta analysis by Arsalidou and Taylor (2011).

#### 4.9.4. ROI temporal dynamics

We explored the dynamics in these ROIs by fitting generalized linear models (GLM) using AFNI (Cox, 1996). Analogous to the whole brain GLM analysis (i.e., boxcar analysis), we spatially smoothed the signal and censored outliers from the regression. In this case, in contrast to the whole brain analysis GLMs, we modeled the trial time using a Finite Impulse Response (FIR) approach, in which each trial was modeled using 17 simple basis functions (tents; Fig 1).

This approach allowed us to take advantage of the short TRs (0.8s) used for the functional acquisition sequence, which were possible due to the ultra-high-field MRI used in the experiment. Modeling the BOLD signal using FIR allowed us to obtain 17 beta estimates *β_F IR_* for each voxel for each of the conditions considered. Note that these estimates model the hemodynamic response directly and, therefore, they do not factor in the lag of the BOLD signal. In order to link each *β_F IR_* to a time in the task, we assumed a lag of 5 seconds in the hemodynamic response.

We obtained a 2*×*2 *β_F IR_*-estimates for the factors TCC (high and low) and satisfiability (satisfiable and unsatisfiable). We explored the dynamics of each ROI by estimating the average *β_F IR_* over all of the voxels from each ROI for each condition. The ROI signal aggregation was performed using python 3.7 and the nilearn library.

#### 4.9.5. Connectivity analysis

Connectivity analysis was performed over the three ROIs. To remove non-neural sources from the neural signal, the motion parameters were regressed out before extracting the relevant ROI signals. We then performed connectivity analysis using two separate approaches.

##### Psychophysiological interaction (PPI)

We performed generalized PPI analyses using AFNI. We ran two separate regressions for each ROI; one for satisfiability and one for TCC. Each PPI regression was estimated according to the following:

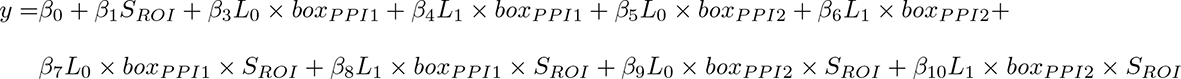

where *L*_0_ corresponds to low TCC (or satisfiable) condition and *L*_1_ corresponds to high TCC (or unsatisfiable) condition. *S_ROI_* is the neural signal of the seed region and *box_i_* corresponds to a boxcar function that separates the items and solving stages, together, into two boxcar functions (PPI-1 and PPI-2) of the same duration (12.5s each; Fig 6). Note that these boxcar functions are different in duration to the ones used for the boxcar GLM analysis. The contrasts of interest (*β*_8_, *β*_7_, *β*_10_ and *β*_9_) captured the PPI effects; that is, the task-dependent connectivity to the ROIs for each of the two periods considered. Additionally, we tested whether there were regions that showed a differential connectivity to an ROI between conditions (i.e., high vs. low TCC, unsatisfiable vs. satisfiable). Explicitly, we performed group level analysis using mixed effects multilevel modeling (Chen et al., 2012) on the contrasts corresponding to *L*_1_ *− L*_0_ (*β*_8_ *− β*_7_ and *β*_10_ *− β*_9_). Results are reported with a clusterwise threshold of *p <* 0.05 corrected for multiple comparisons across the whole brain, using an uncorrected voxelwise threshold of *p <* 0.001.

It is worth noting that the interaction between box-car functions and the seed region (*box × S*) was estimated via deconvolution. That is, the BOLD time series of each seed region was deconvolved with a canonical HRF (AFNI: *BLOCK(0.1,1)*) and then multiplied with the psychological boxcar function. This was convolved back with the same HRF to form a predicted PPI time series at the hemodynamic response level (BOLD), at which the regression takes place.

##### Granger causality

Additionally, we performed Granger Causality (GC) analysis on the three ROIs. To do this, we first fitted a DCM to the BOLD time series of these ROIs. This was done to ensure that the DCM captured all the task-relevant events and controls not strictly related to the internal solving process itself (e.g., onset of decision screen). We report the exact specification of the DCM in Appendix A.2. We then extracted the residual series of the DCM model for each region. We refrained from deconvolving the BOLD residuals (in accordance with Seth et al. (2013)) because deconvolution is a smoothing operation that introduces spurious lead-lag relationships.

GC emerges when lagged outcomes of a variable *correlate* significantly with values of another variable. As such, GC is closely linked to *cross-autocorrelations*. Typically, GC is analyzed in the context of a Vector Auto Regression (VAR), i.e., a model whereby a vector of outcomes is driven by a finite number of lags of itself. GC emerges when the presence of lags of one variable significantly improves the fit (maximum likelihood value) of another variable. If this is the case, the former “Granger causes” (GCs) the latter. We ran a VAR on the error series augmented with the error series during the solving stage only, and determined incremental GC of one series on another during problem-solving.

In order to reach a GC statistic at the group level we carried out the following procedure. We first ran a VAR for each subject. Each subject’s VAR maximum lag was determined by comparing AIC (Akaike Information Criterium) for lags up to 10. From each regression we extracted 5 GC statistics for each ROI: 2 GCs from lagged time series of each of the other two ROIs and 3 GCs (one for each ROI) from the lagged time series of the solving stage. This process generates 15 GC statistics per subject. To correct for multiple comparisons among these we performed standard Bonferroni correction.

To determine statistical significance at the group level, a standard binomial test was then employed to determine the significance of the frequency of rejections (of no GC) across the 20 participants. A *p* level of was 0.05 used. FWE correction was applied using Holm-Bonferroni correction over the 15 tests.^3^

The Matlab method gctest was used to implement the Granger Causality estimations.

#### 4.10. Data and code availability

The data analysis code and the behavioral data will be made available upon publication at the Open Science Framework (OSF). The software for the knapsack decision task will be made available there as well. The anonymized neuroimaging data will be made available (in BIDS format) upon publication. The software for the knapsack optimization task and mental arithmetic task correspond to those employed by Franco et al. (2021b) and are available at the OSF (DOI 10.17605/OSF.IO/T2JV7).

### Appendix A fMRI preprocessing

#### A.1 General pipeline

Raw images were organized and converted to the relevant format according to the (BIDS) standards.

Pulse and cardiac noise were regressed out from the functional scans using RETROICOR. These were then slice-time corrected and the volumes were motion-corrected by registering to the first volume of the first functional run. The anatomical (T1) image was down-sampled to the functional EPI resolution (1.6*mm*^3^) and the mean BOLD volume of the first run was co-registered to the down-sampled anatomical scan. This transformation was applied to all of the BOLD volumes. Afterwards, each participant’s anatomical scan was used for calculation of transformation parameters to normalize the functional images into the Montreal Neurological Institute (MNI) space (Fig 7).

**Figure 7:**
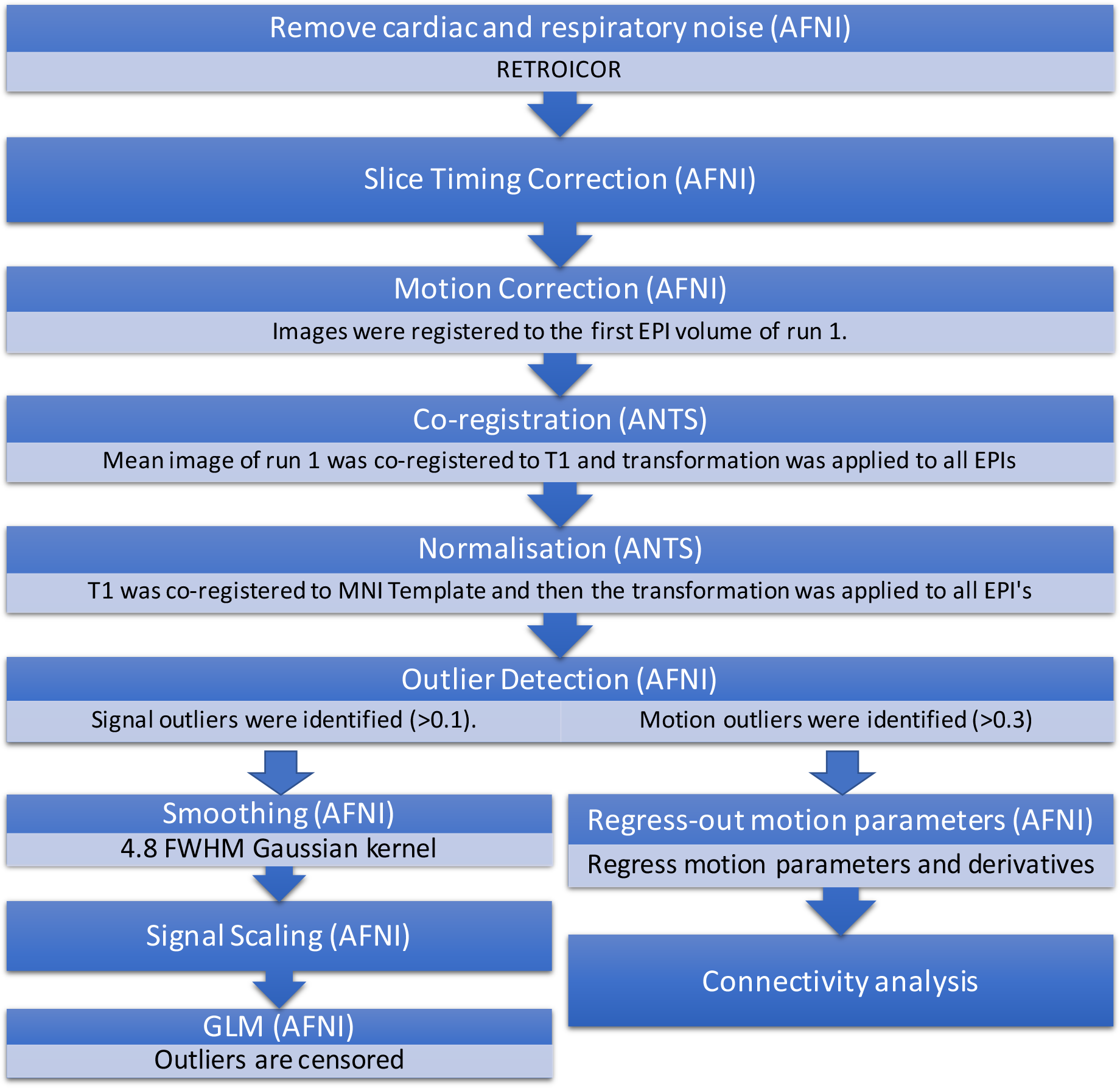
fMRI data preprocessing pipeline. Depiction of the preprocessing steps used prior to the statistical analyses performed on the functional data. The preprocessing steps, up to outlier detection, are shared across all types of analysis. Afterwards, preprocessing steps differ between GLMs and functional connectivity models.

Whole-brain analyses were performed by fitting generalized linear models (GLM) using AFNI (Cox, 1996). Before the regressions were implemented, we spatially smoothed the functional volumes with a 4.8mm FWHM Gaussian kernel. Each voxel’s signal was then scaled (per run) to have the same mean (100). Additionally, volumes with motion or signal outliers were censored from each of the regressions. Regressions were performed using the 3dREMLfit algorithm in AFNI. Group level statistical tests were performed using mixed effects multilevel modeling (Chen et al., 2012).

For the connectivity analyses the volumes were not smoothed, but motion parameters were regressed out before extracting the relevant ROI signals. Whitened (ARMA(1,1)) residuals were used in the subsequent analyses.

#### A.2 Granger Causality preprocessing: DCM

Additional to the preprocessing steps presented in the previous section we fitted a dynamic causal model (DCM) before Granger causality (GC) analysis. This was done in order to remove signals of no interest related to perceptual processes related to screen and stage changes. The model was fit using the SPM12 software (https://www.fil.ion.ucl.ac.uk/spm/software/spm12/) and the residuals of the resulting model were then used to fit the VAR model and test for GC (see section 4.9.5). In this section we describe the DCM used.

Let *z* denote a vector of neural activity in 3 regions, indexed *i* (= *rAI, rAG, dACC*).

Let *v_j_* denote conditions; they reflect stages in the task conditional on properties of the instance (i.e., satisfiability and TCC). The variable is a dummy variable that takes the value of 1 when the *j* condition is ON the screen:

- *j* = 1: items stage and solving stage (25s),
- *j* = 2: response stage (2s),

Additionally, let *o_j_* denote onsets of conditions; that is, when a stage becomes visible on the screen. We follow the SPM’s notation to describe the model employing three different types of matrices. Matrix

A specifies the baseline effective connectivity. Matrix *B*^(^*^j^*^)^ denotes the modulation of effective connectivity due to experimental condition *j*. Finally, *C*^(^*^j^*^)^ captures the change of the neural response due to the onset of condition *j*.

The DCM fit is described by the following three equations; one for each ROI: For rAG:

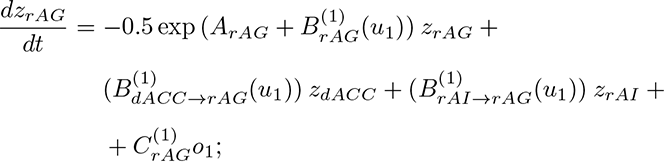

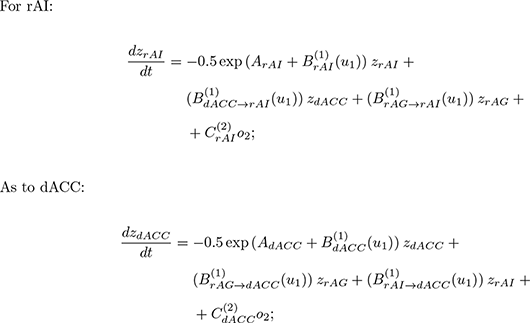

It is worth noting the asymmetries between regions in our specification. These are found in the burst of activity in the model (C matrix). Specifically, we expected the AG to be a processing unit with activity starting quickly from the items stage in the task; this is reflected in the *C_rAG,_*_1_*o*_1_ term in the rAG equation. In contrast, we expected the AI and dACC to present burst activity related to control and monitoring signals at the moment the solving stage ends (i.e., *C_rAI,_*_2_*o*_2_ and *C_dACC,_*_2_*o*_2_). Besides this asymmetry, the model allows for a symmetric inter-connectivity between ROIs during the items and solving stage of the task.

### Appendix B Tables and Figures

**Table 4:**
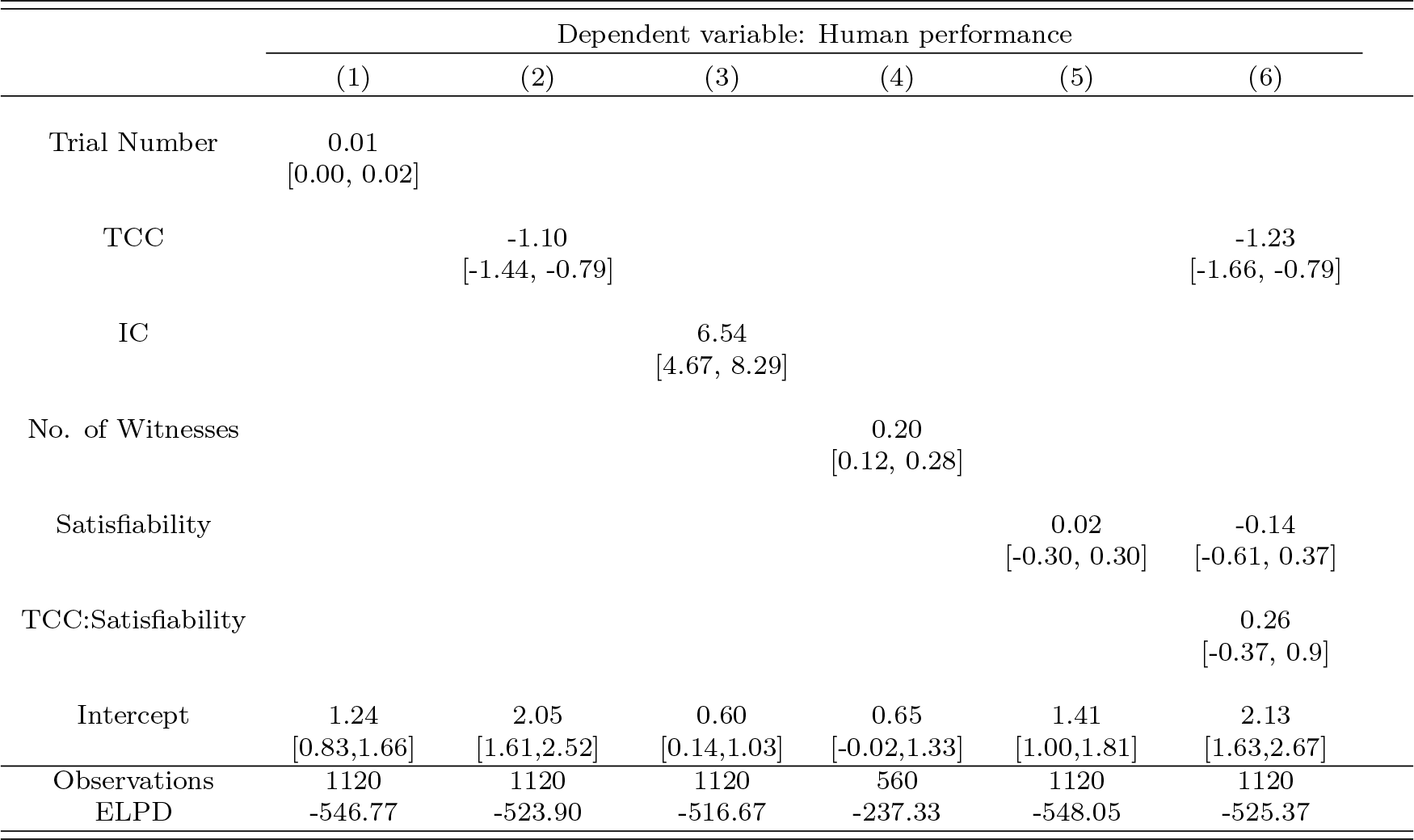
Human performance in the knapsack decision task. Logistic regressions with random intercept effects for participants relating the accuracy on an instance and trial number (1), typical-case complexity (TCC) (2), instance complexity (IC) (3), the number of witnesses (4), satisfiability (5), as well as TCC and satisfiability (6). Parameter estimates correspond to the median of the posterior distribution (β0.5) and the 95% HDI credible interval (HDI0.95). ELPD denotes the expected log posterior predictive density.

|  | Dependent variable: Human performance |  |  |  |  |  |
| --- | --- | --- | --- | --- | --- | --- |
|  | (1) | (2) | (3) | (4) | (5) | (6) |
| Trial Number | 0.01<br>[0.00, 0.02] |  |  |  |  |  |
| TCC |  | -1.10<br>[-1.44, -0.79] |  |  |  | -1.23<br>[-1.66, -0.79] |
| IC |  |  | 6.54<br>[4.67, 8.29] |  |  |  |
| No. of Witnesses |  |  |  | 0.20<br>[0.12, 0.28] |  |  |
| Satisfiability |  |  |  |  | 0.02<br>[-0.30, 0.30] | -0.14<br>[-0.61, 0.37] |
| TCC:Satisfiability |  |  |  |  |  | 0.26<br>[-0.37, 0.9] |
| Intercept | 1.24<br>[0.83,1.66] | 2.05<br>[1.61,2.52] | 0.60<br>[0.14,1.03] | 0.65<br>[-0.02,1.33] | 1.41<br>[1.00,1.81] | 2.13<br>[1.63,2.67] |
| Observations | 1120 | 1120 | 1120 | 560 | 1120 | 1120 |
| ELPD | -546.77 | -523.90 | -516.67 | -237.33 | -548.05 | -525.37 |

**Figure 8:**
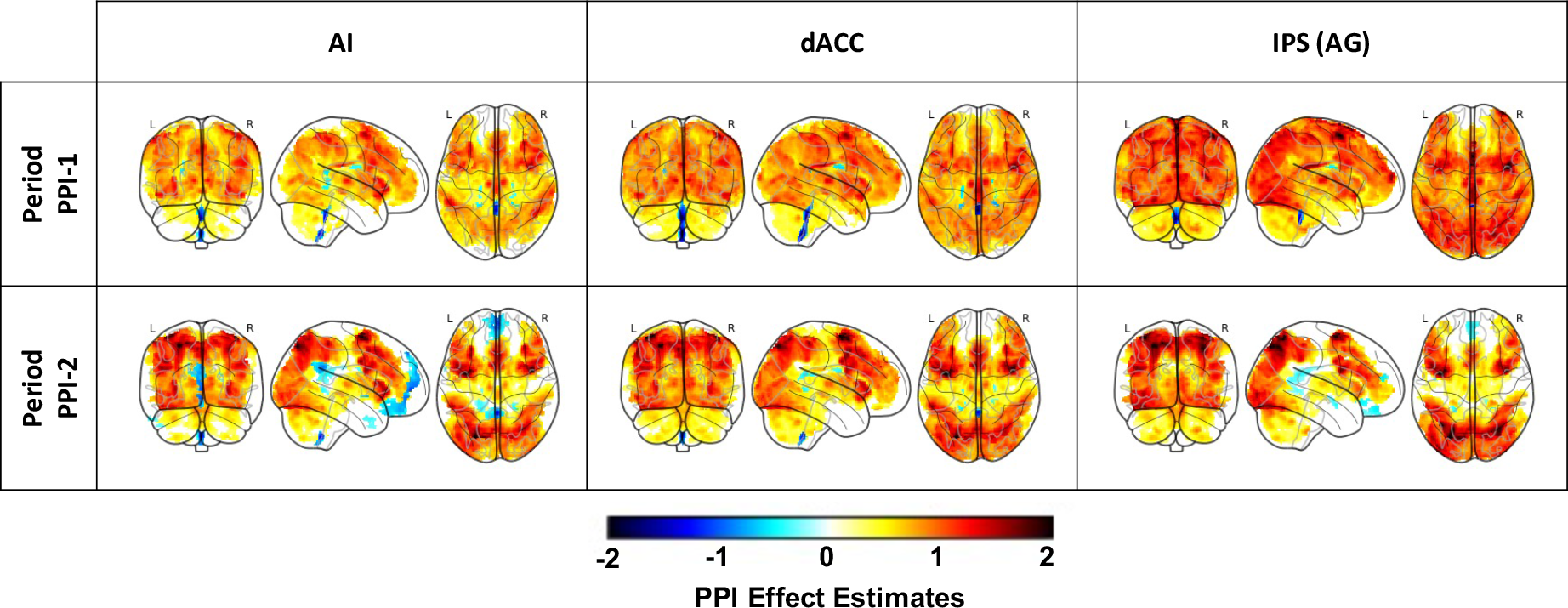
PPI supplementary results. The effect of the task on the connectivity to each of the three seed regions used for the analysis (dACC, rAG and rAI). Each column shows the PPI effect for a different seed region. Each row displays the period of the task considered. Activation patterns represent the significant effect estimates for the PPI on instances with low TCC. The effect for instances with high TCC is not displayed, but the difference with respect to the activation signatures shown here are small. Indeed, the only significant differences between conditions are presented in figure 6a. *Significant cluster-wise FWE-corrected (p <* 0.05*) clusters (with an uncorrected threshold of p <* 0.001*) are presented*.

### Appendix C Replication of previous behavioral results

#### C.1 Knapsack decision task

An additional aim for this study was to reproduce the key findings presented by Franco et al. (2021b). We first looked at the effect of experience on accuracy and found a non-significant improvement as the task progressed (*β*_0.5_ = 0.009, *HDI*_0.95_ = [*−*0.001, 0.021], main effect of trial number on performance, generalized logistic mixed model (GLMM); Table 4 Model 1). This marginal improvement in the task performance might seem to contradict previous results, which suggest that neither experience with the task nor mental fatigue affected task performance. However, unlike Franco et al. (2021b), we performed the task in the scanner, thus this discrepancy could be due to acclimatization to the scanner.

In the main text we show that the results regarding TCC and satisfiability are mirrored by our data and statistical analyses. Specifically, our findings corroborate the significant effect of TCC on performance and replicate a null effect of satisfiability on performance. Additionally, other key findings in their study were related to two *solution-space* metrics of complexity: The number of solution witnesses and instance complexity (IC). The former is defined as the number subsets of items that satisfy both profit and capacity constraints while IC is defined as the distance between the level of the profit constraint (target profit) and the maximum value attainable in the corresponding instance of the optimization variant of the 0-1 knapsack problem. Specifically,

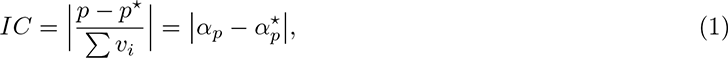

where *p* is the target profit of the decision instance and *p^*^* is the maximum value achievable in the corresponding optimization instance, that is, the maximum value that can be packed into the knapsack given the same set of items *I* and the same capacity constraint *c*. *α_p_* and *α^*^* denote the normalized values of target profit and optimum value, respectively.

In order to estimate these metrics, unlike TCC, the problem needs to be solved. Concretely, harder versions of the problem needs to be solved. For IC to be estimated, the optimization variant of the knapsack problem needs to be solved, while for the number of witnesses all of the possible sets of items that satisfy the constraints need to be found. This makes estimating these metrics more computationally intensive than estimation of TCC. Despite this drawback, these metrics capture the hardness of a single instance of the problem and therefore are more precise when predicting performance for each instance compared to TCC, which captures the average hardness of an ensemble of random instances.

Franco et al. (2021b) showed that human performance was affected by both IC and the number of witnesses. Here we reproduced these findings. We found that higher values of IC were related to higher accuracy (*β*_0.5_ = 6.54, *HDI*_0.95_ = [4.67, 8.29], main effect of IC, GLMM; Table 4 Model 3; Fig 9). Similarly, among satisfiable instances, we found that a higher number of witnesses was related to better performance (*β*_0.5_ = 0.20, *HDI*_0.95_ = [0.12, 0.28], main effect of number of witnesses in satisfiable instances, GLMM; Table 4 Model 4). It is worth noting that the number of witnesses can only explain variability among satisfiable instances since all unsatisfiable instances have 0 witnesses.

**Figure 9:**
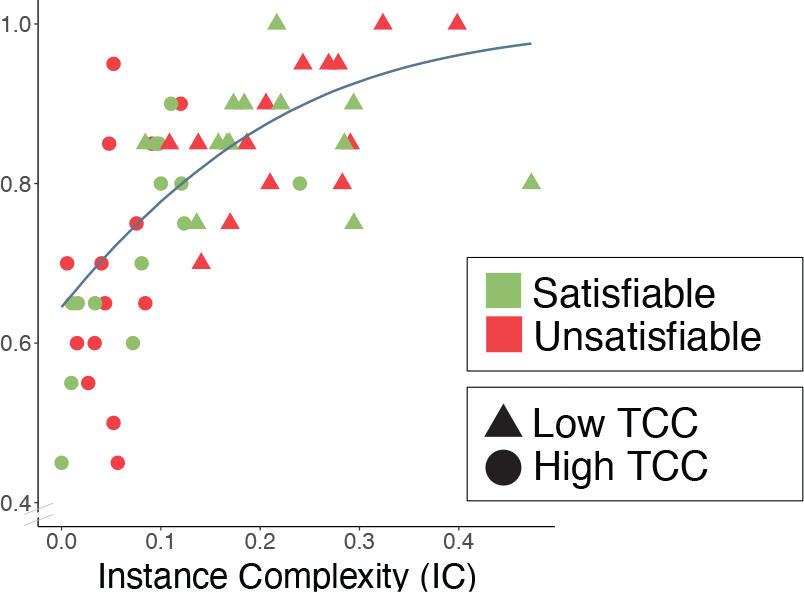
Relation between IC and human performance in the knapsack decision task. Mean accuracy per instance and the marginal effect of IC on human performance (GLMM; Table 4 Model 3). Higher IC is related to lower computational hardness. Instances are categorized by their TCC (shape) and satisfiability (color).

Overall, these results replicate previous findings (Franco et al., 2021b) and validate that the experimentally modulated variable (TCC) successfully varied the computational difficulty of the task.

#### C.2 Knapsack optimization task

In this task, participants were asked to solve a number of instances of the (0-1) knapsack optimization problem (Fig 10(a)). In each trial, they were shown a set of items with different weights and values as well as a capacity constraint. Participants had to find the subset of items that maximized total value subject to the capacity constraint. This means that while in the knapsack decision task, participants only needed to determine whether a solution existed, in the knapsack optimization task, they also needed to determine the nature of the solutions (i.e., the items in the optimal knapsack).

**Figure 10:**
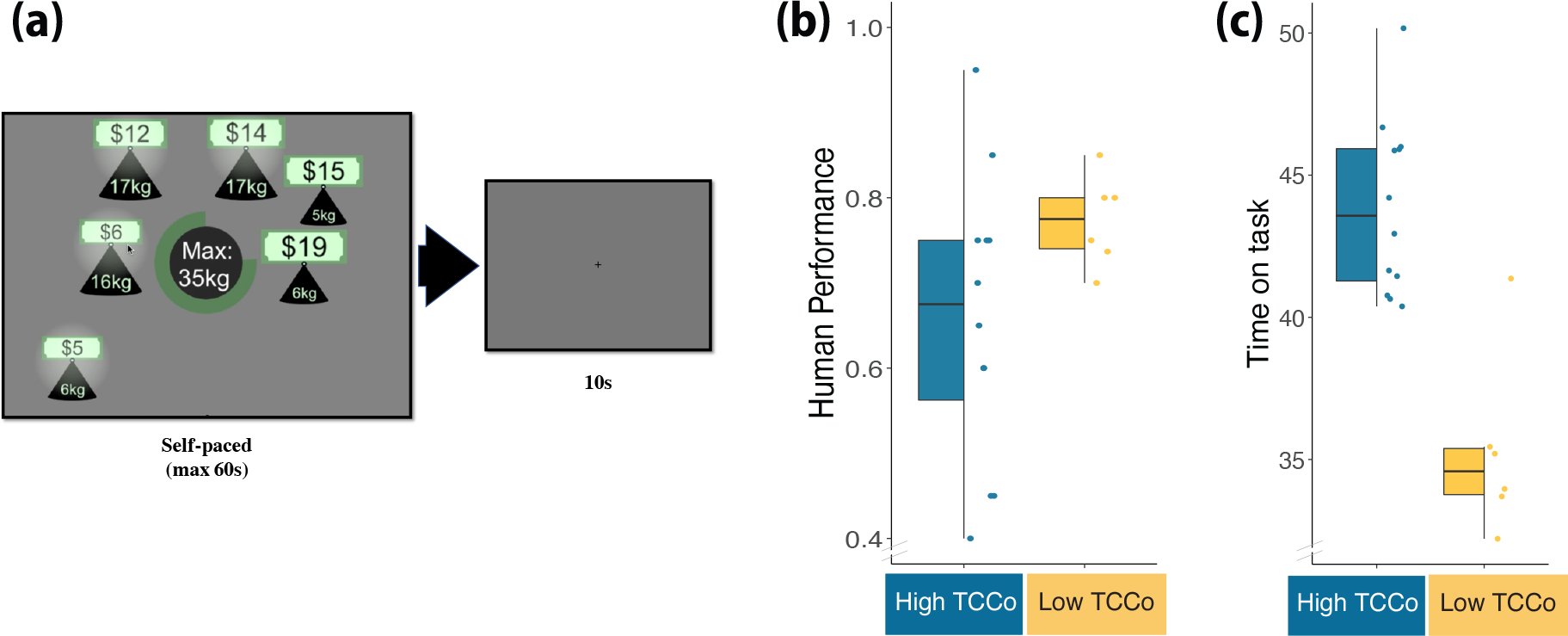
Knapsack optimization task. (a) Experimental design. Participants were presented with a set of items of different values and weights together with a capacity constraint shown at the center of the screen. The green circle at the center of the screen indicated the time remaining in this stage of the trial. Participants had to find the subset of items with the highest total value subject to the capacity constraint. This stage lasted up to 60 seconds. Participants selected items by clicking on them and had the option of submitting their solution before the time limit was reached. After the time limit was reached or they submitted their solution, a fixation cross was shown for 10 seconds before the next trial started. **(b) TCC***_O_* **and human performance.** Human performance corresponds to mean computational performance on each instance. **(c) TCC***_O_* **and time-on-task.** Mean time spent before skipping to the response screen. *Each dot represents an instance and is categorized according to its TCC_O_. The box-plots represent the median, the interquartile range (IQR) and the whiskers extend to a maximum length of 1.5*IQR*

For this task we aimed at replicating the results found by Franco et al. (2021b). In their study, a metric of complexity (TCC*_O_*) was introduced as an extension of the TCC metric to optimization problems. Specifically, TCC*_O_* was defined as the TCC of the decision of determining whether the optimal profit (*α^∗^*) is attainable given the capacity constraint. We expected to replicate the negative effect of TCC*_O_* on performance and its positive effect on time-on-task.

The task consisted of a single solving stage (60 seconds) and an inter-trial interval (fixation cross for 10 seconds). During the solving stage the items and the capacity constraint were presented in the same way as in the knapsack decision task. Unlike in the decision task, however, there was no target profit and participants were able to add and remove items to/from the knapsack by clicking on the items. An item added to the knapsack was indicated by a halo around it (Fig 10). Participants could submit their solution before the time limit was reached. If participants did not submit within the time limit, the items selected at the end of the trial were automatically submitted as the solution. Participants were then shown a fixation cross (10 seconds) before the start of the next trial.

Each participant completed 18 trials (2 blocks of 9 trials with a rest period of 60 seconds between blocks). Each trial presented a different instance of the knapsack optimization problem with varying levels of computational complexity. Specifically, we employed the same instances of the knapsack optimization problem used in Franco et al. (2021b). In their study, 12 instances were selected to have high TCC*_O_* and 6 instances were selected to have low TCC*_O_* . All instances had *N* = 6 items and *w_i_*, *v_i_*, *c* and *p* were integers.

The order of presentation of instances in the task was randomized for each participant. For the analysis, we excluded 1 trial from one participant because solutions were submitted after less than 1 second into the task. Additionally, 2 participants were excluded from the analysis of time-on-task because they never submitted a solution before time ran out.

We investigated two variables related to behavior: performance and time-on-task. Performance was quantified by *computational performance*, which captures participants’ ability to find the optimal solution. Specifically, it is defined as a binary variable that is equal to 1 if the participant obtained a value equal to the maximum value obtainable in the instance, and 0 otherwise. It is worth noting that an instance was only characterized as correct if the sum of weights did not exceed the capacity constraint. Mean computational performance was 69.6% (min = 0.18, max = 1, *SD* = 0.25) and the capacity constraint was only violated in 3.9% of instances. Additionally, we investigated time-on-task. In contrast to the decision variant, the optimization task was self-paced and, as such, participants were allowed to submit their answer before the time limit (60s) was reached. We recorded the time participants spent in the solving stage before submitting their candidate solution. Participants spent on average 41.0 seconds on an instance (min = 21.0, max = 55.8, *SD* = 8.1).

We first replicated the effect of trial number in the task. We found that performance did not change throughout the task (*β*_0.5_ = 0.03, *HDI*_0.95_ = [*−*0.02, 0.09], main effect of trial number on computational performance, GLMM; Table 5 Model 1), nor did the time-on-task per instance (*β*_0.5_ = *−*0.02, *HDI*_0.95_ = [*−*0.03, 0.22], main effect of trial number on time-on-task, CLMM; Table 5 Model 3). These results suggest, in line with previous results (Franco et al., 2021b), that neither experience with the task nor mental fatigue affected the quality and speed of finding the a solution.

**Table 5:** Computational performance and time-on-task in the knapsack optimization task. Models on computational performance represent logistic regressions with random intercept effects for participants. Regression parameters relate performance to trial number (1), and optimization typical-case complexity (TCC_O_ ) (2). Models on time-on-task represent censored linear regressions (with random intercept effects for participants) relating time spent on an instance to trial number (3), and optimization typical-case complexity (TCC_O_ ) (4). Parameter estimates correspond to the median of the posterior distribution (β0.5) and the 95% HDI credible interval (HDI0.95). ELPD denotes the expected log posterior predictive density.

|  | Dependent variable |  |  |  |
| --- | --- | --- | --- | --- |
|  | Computational performance |  | Time-on-task |  |
|  | (1) | (2) | (3) | (4) |
| Trial Number | 0.03<br>[-0.02,0.09] |  | -0.02<br>[-0.26,0.22] |  |
| $TCC_O$ | | -0.75<br>[-1.35,-0.14] | | 8.78<br>[6.45,10.97] |
| Intercept | 0.84<br>[0.05,1.7] | 1.61<br>[0.74,2.45] | 41.21<br>[36.51,45.47] | 35.12<br>[30.98,39.98] |
| Observations | 359 | 359 | 323 | 323 |
| ELPD | -184.19 | -181.54 | -1228.64 | -1199.62 |

Finally, we studied the effect of TCC*_O_*. We expected that performance in instances with *high TCC_O_* (instances whose solutions have a corresponding decision problem with high TCC) would be lower than in instances with *low TCC_O_* (instances whose solutions have a corresponding decision problem with low TCC). We, indeed find this effect on both computational performance and time-on task. Mean computational performance was lower in instances with high TCC*_O_* , relative to those with low TCC*_O_* (*β*_0.5_ = *−*0.75, *HDI*_0.95_ = [*−*1.35*, −*0.14], main effect of TCC*_O_* on performance, GLMM; Fig 10b; Table 5 Model 2). Similarly, we found a positive effect of TCC*_O_* on time-on-task (*β*_0.5_ = 8.78, *HDI*_0.95_ = [6.45, 10.97], main effect of TCC*_O_* on time-on-task, CLMM; Fig 10c; Table 5 Model 4). These results replicate those found by Franco et al. (2021b).

### Appendix D Cognitive function tasks

In a previous study we tested participants’ performance on five aspects of cognitive function that we considered relevant for the knapsack tasks (Franco et al., 2021b). Explicitly, we assessed working memory, episodic memory, strategy use, processing and psychomotor speed, as well as mental arithmetic. We were interested in finding links between these cognitive capacities and the ability to solve the knapsack task. A complex task that would arguably require the deployment of these other, more basic, cognitive abilities. Our original study lacked the power to identify reliably correlations between performance in these cognitive tasks and performance in the knapsack tasks.

In this study we tested participants on the same five aspects of cognitive function with the aim of increasing the power of these exploratory tests. For this purpose, we aggregated the data collected in this study with that collected by (Franco et al., 2021b) and estimated the same correlations presented in our previous study.

Following the approach by Franco et al. (2021b) we administered a set of tasks from the Cambridge Neuropsychological Test Automated Battery (CANTAB; Cognition, 2017). Specifically, we asked participants to perform the Paired Associates Learning (PAL), Spatial Working Memory (SWM) and Spatial Span (SSP). Additionally, participants solved a set of mental arithmetic problems (Cappelletti et al., 2001). Below we describe each of the tests performed:

#### Paired Associates Learning (PAL)

Boxes are displayed on the screen and open one by one in a randomized order to reveal patterns hidden inside. The patterns are then displayed in the middle of the screen, one at a time, and the subject must touch the box where the pattern was originally located.

#### Spatial Working Memory (SWM)

The test begins with colored boxes being shown on the screen. The aim of this test is that, by touching the boxes and using a process of elimination, the subject should find one ‘token’ in each of the boxes and use them to fill up an empty column on the right hand side of the screen. The computer will never hide a token in the same colored box, so once a token is found in a box the participant should not return to that box to look for another token.

#### Spatial Span Task (SSP)

White squares briefly change color in a variable sequence. The participant must remember the sequence and then touch the squares in that same order. The sequence length increases through the test. There are up to 3 attempts at each sequence length and the test terminates if all three are failed.

#### Mental Arithmetic Task

Participants were asked to answer a set of 33 mental arithmetic problems. They were given 13 seconds to solve each problem. The task involved addition and division of numbers, as well as questions in which they were asked to round to the nearest integer the result of an addition or division operation.

From performance in these tasks we estimated five metrics of cognitive capacities and estimated their correlation with participant’s performance on the knapsack decision and optimization tasks. Results are presented in Table 6. We found, after correcting for multiple comparisons using Holm-Bonferroni correction, a significant positive effect between performance in the knapsack optimization task and performance in the mental arithmetic task (*ρ* = 0.617 at FWE-corrected *α* = 0.05). Additionally, we found (at FWE-corrected *α* = 0.10) a negative correlation between the *strategy use* metric and performance in the knapsack decision task (*ρ* = *−*0.421). The SWMS metric encodes the number of times a subject begins a new search pattern from the same box they started with previously in the SWM task. Therefore, a lower score is interpreted as higher strategy use (1 = they always begin the search from the same box). These results suggest that participants that use a planned strategy in SWM perform better in the knapsack decision task.

**Table 6:** Pearson correlations between performance in the knapsack tasks and cognitive abilities. Performance in the knapsack decision task is characterized by accuracy and in the knapsack optimization task is characterized by computational performance. The cognitive abilities measured used were mental arithmetic, episodic memory (PALFAMS28), working memory (SSPFSL), strategy use (SWMS) and spatial working memory (weighted SWMTE, with errors on easier tasks being weighted more). P-values are shown without multiple comparisons correction. ^1^In the mental arithmetic task *df* = 37. *Note: FWE significance ^∗^ <0.1; ^∗∗^ <0.05; ^∗∗∗^ <0.01 is assessed employing Holm-Bonferroni correction*.

| Task | Knapsack<br>decision | Knapsack<br>optimization |
| --- | --- | --- |
| Mental<br>arithmetic | 0.311<br>(0.156)<br>p=0.054 | 0.617***<br>(0.129)<br>p=0.000 |
| Episodic<br>memory | 0.098<br>(0.161)<br>p=0.549 | -0.017<br>(0.162)<br>p=0.918 |
| Working<br>memory | 0.033<br>(0.162)<br>p=0.839 | 0.218<br>(0.158)<br>p=0.177 |
| Strategy<br>use | -0.421*<br>(0.147)<br>p=0.007 | -0.336<br>(0.153)<br>p=0.034 |
| Spatial working<br>memory | -0.360<br>(0.151)<br>p=0.023 | -0.348<br>(0.152)<br>p=0.028 |
| Degrees of<br>freedom<sup>1</sup> | 38 | 38 |

### Appendix E Neural correlates and dynamics of accuracy

#### E.1 Neural correlates

It has been hypothesized that FPN as well as CON regions encode task signals related to error detection and error expectation (Neta et al., 2017, 2014; Dosenbach et al., 2006). Although participants did not receive any feedback during the task, we expected to see error related signals during later stages of the trial. Although these signals would not represent the integration of novel exogenous information (since there was no feedback) we conjectured that participants would represent a subjective belief on the expected accuracy (or reward) of their answer (e.g, Duverne and Koechlin, 2017).

We found only one significant cluster during the solving stage (in period one) (Fig 11a; Table 7). The other significant clusters were identified during the response stage (Fig 11b; Table 7). In line with out hypothesis, during the response stage we found that activity in both the FPN and CON was positively correlated with erring. Specifically, a higher activity was found for incorrect trials in the AI (bilaterally), dACC, left MFG and the right inferior frontal gyrus. Additionally to these regions, which are commonly associated with the MDS, we also found significant activation in the SFG (bilaterally), ACC and paracingulate gyrus.

**Figure 11:**
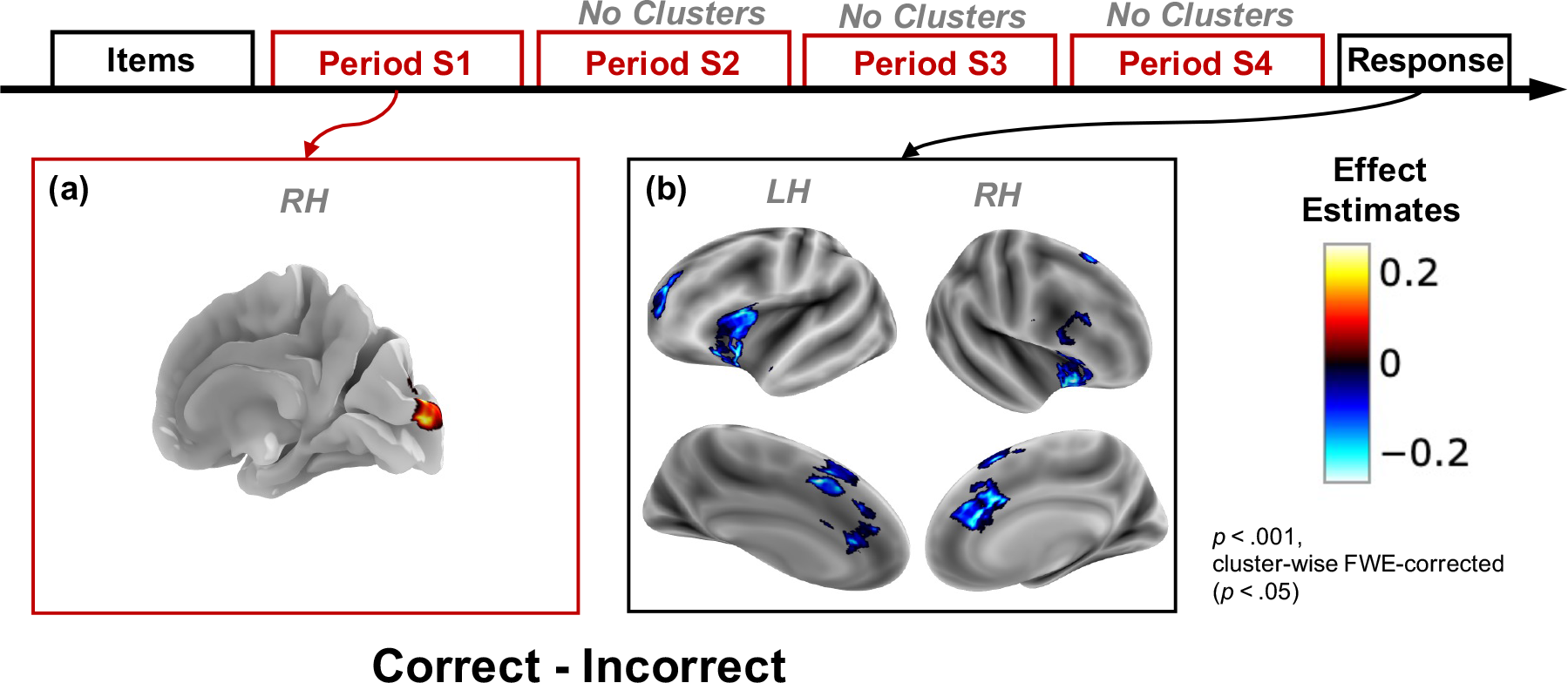
Neural correlates of accuracy. Brain activation effect estimates (*β*) for the correct vs. incorrect contrast (*βcorrect − βincorrect*). A positive contrast represents a higher BOLD activity on instances that were answered correctly. Significant cluster-wise FWE-corrected (*p <* 0.05) clusters (with an uncorrected threshold of *p <* 0.001) are presented for each of the contrasts estimated using the Boxcar analysis. Each panel represents a different period in the trial. **(a)** Period S1, **(b)** response stage. No significant clusters were found for the contrasts during periods S2-S4 of the solving stage.

**Table 7:** Accuracy clusters. . Significant cluster-wise FWE-corrected (p < 0.05) clusters (using an uncorrected threshold of *<* 0.001) from the *Correct-Incorrect* contrast. Coordinates are in MNI space.

| Stage | Region | Side | Cluster statistics |  |  | Peak statistics |  |  |  |
| --- | --- | --- | --- | --- | --- | --- | --- | --- | --- |
| | | | Volume($mm^3$) | $\beta_{mean}$ | SEM | $\beta_{peak}$ | x | y | z |
| S1 | Occipital cortex | RH | 569.3 | 0.21 | 0.003 | 0.31 | 10 | -100 | 8 |
| Response | AI | RH | 2146.3 | -0.22 | 0.002 | -0.33 | 30 | 23 | -8 |
|  | AI | LH | 2048.0 | -0.26 | 0.002 | -0.40 | -48 | 18 | -12 |
|  | dACC | LH | 1953.8 | -0.24 | 0.002 | -0.38 | -2 | 22 | 40 |
|  | SFG | RH | 1007.6 | -0.18 | 0.003 | -0.28 | 2 | 23 | 60 |
|  | MFG | LH | 974.8 | -0.19 | 0.002 | -0.27 | -27 | 50 | 16 |
|  | SFG | LH | 684.0 | -0.18 | 0.003 | -0.26 | -2 | 10 | 60 |
|  | Inferior frontal gyrus | RH | 602.1 | -0.22 | 0.003 | -0.29 | 50 | 17 | 31 |
|  | ACC | LH | 520.2 | -0.18 | 0.002 | -0.25 | -3 | 31 | 26 |
|  | Paracingulate gyrus | LH | 491.5 | -0.24 | 0.004 | -0.31 | -5 | 9 | 50 |

The results of this analysis confirm our hypothesis by outlining a set of regions in both FPN and CON that encode errors during the response stage. In contrast, this analysis did not result in any other clusters during the solving stage that correlated negatively with accuracy. The lack of negative correlation during the late periods of the solving stage could be due to variability in the signal during the solving stage. Indeed, during this stage participants might be updating their accuracy expectation as well as their candidate response. Since our accuracy contrast is based on the answer provided during the response stage, it stands to reason that our analysis does not capture accuracy markers during the solving stage because we do not have a measure of accuracy during this period. It is worth noting that we found one significant cluster during the solving stage (in period one) that correlated positively with accuracy in the occipital cortex. This could reflect attentional differences, early in the trial, which affect performance on the trial.

#### E.2 Neural dynamics

We were particularly interested in exploring how the neural correlates of accuracy interact with the proposed metrics of complexity and proof hardness. To do this we studied the dynamics of neural markers of accuracy (employing FIR analysis) for each metric of interest. We first analyzed the interaction effect between correctness and TCC on neural dynamics. We found that for instances with low TCC, there was a significant effect of correctness of the instance from early on in the trial in the IPS. Similarly, midway through the trial a significant accuracy neural marker appeared in the AI for instances with low TCC (Fig 12). This effect was mainly due to a significantly lower BOLD signal on incorrect instances with low TCC. Similarly, when studying the interaction effect between correctness and satisfiability we found a consistent significant effect of accuracy but only during satisfiable instances in the IPS, which was driven, as well, by a lower BOLD activity on incorrect instances (Fig 13). Importantly, this significant contrast showed from the moment the trial started, suggesting that for satisfiable instances the accuracy could be predicted from early on in the trial.

**Figure 12:**
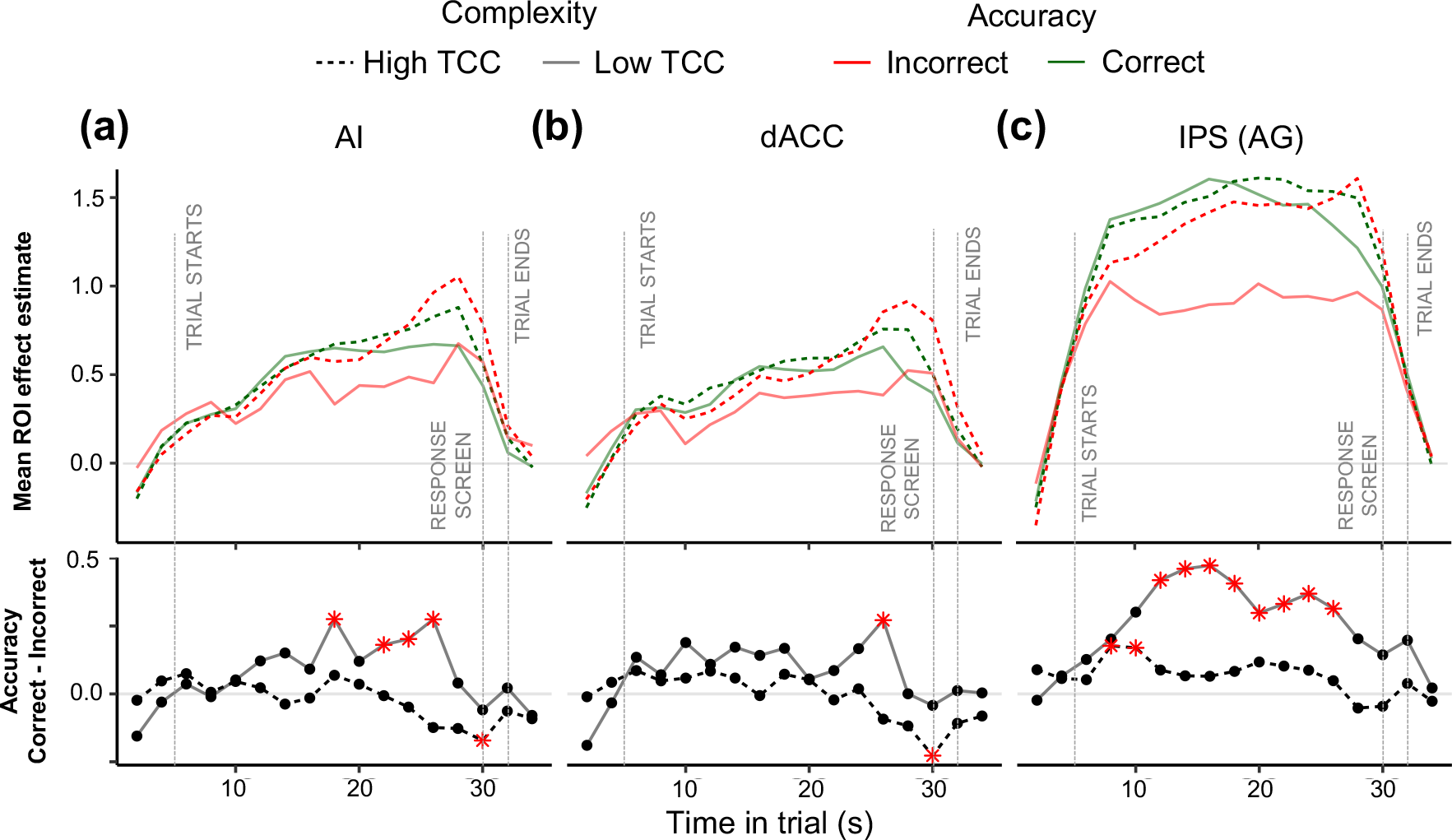
Accuracy and complexity. Mean effect estimate (*β*) of each ROI against time in trial. The effect at each time point represents the mean *β_F IR_* over all of the voxels from each ROI: right AI **(a)**, dACC **(b)**, and right IPS cluster extending to the angular gyrus **(c)**. In the top row of figures the *β_F IR_* ’s characterize the coefficients of an FIR regression with four conditions: accuracy*×*TCC. The *β_F IR_* parameters are aligned to the BOLD signal, which has a lag with respect to the task time. The gray vertical lines represent the task-events assuming a 5 seconds BOLD signal lag. The second row shows the accuracy contrasts (*βcorrect − βincorrect*) for different levels of TCC. Red stars represent significance at a 0.05 significance level.

**Figure 13:**
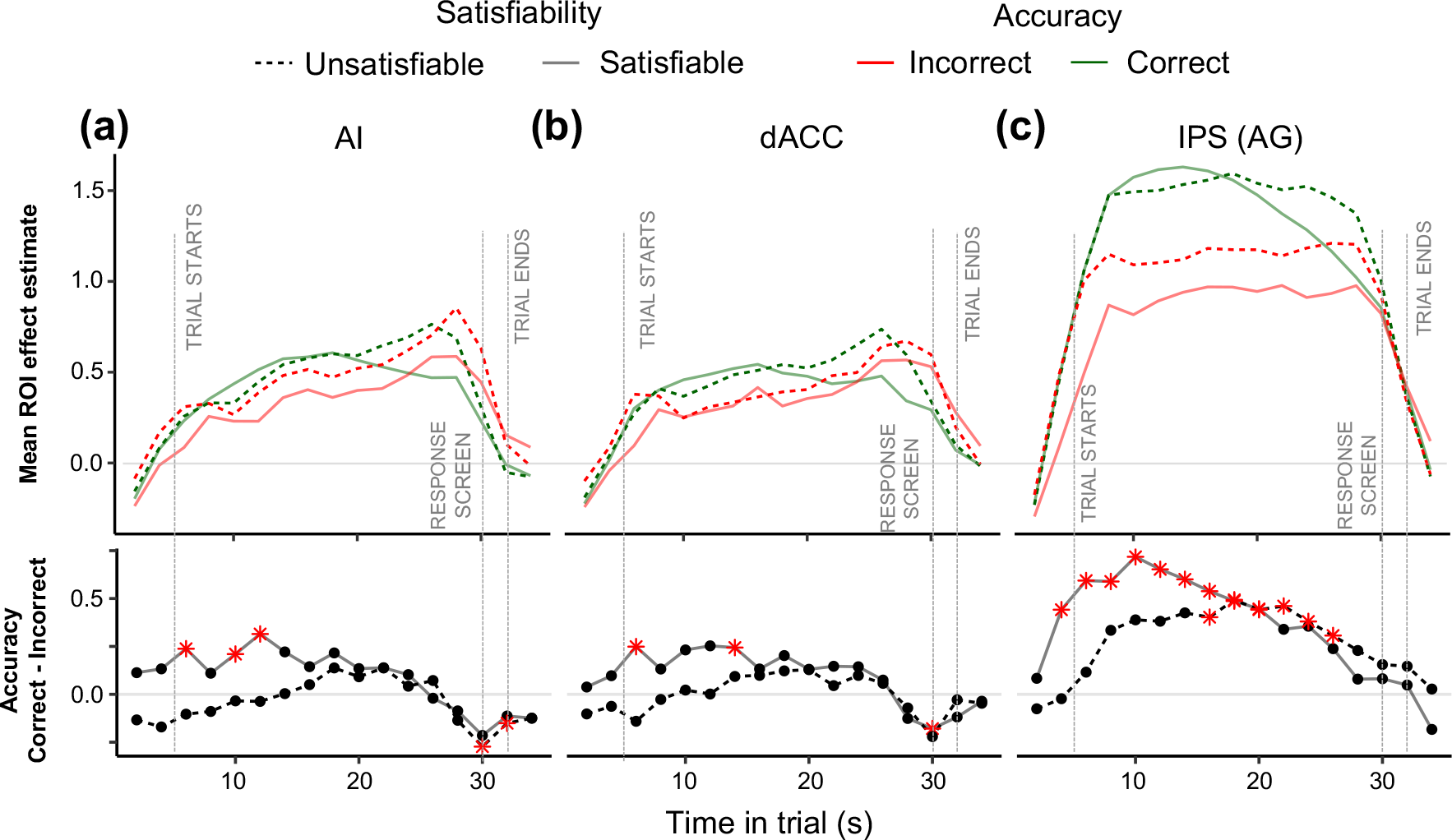
Accuracy and satisfiability. Mean effect estimate (*β*) of each ROI against time in trial. The effect at each time point represents the mean *β_F IR_* over all of the voxels from each ROI: right AI **(a)**, dACC **(b)**, and right IPS cluster extending to the angular gyrus **(c)**. In the top row of figures the *β_F IR_* ’s characterize the coefficients of an FIR regression with four conditions: accuracy*×*satisfiability. The *β_F IR_* parameters are aligned to the BOLD signal, whilst the gray vertical lines represent the task-events. The second row displays the accuracy contrasts (*βcorrect − βincorrect*) separately for satisfiable and unsatisfiable instances. Red stars represent significance at a 0.05 significance level.

Overall, our results suggest a link between neural markers of accuracy and metrics of computational difficulty. This relation was particularly evident in the IPS and marginally in the AI. These findings suggest that neural activity in the IPS is associated with accuracy, especially on instances with low computational difficulty (i.e., low proff hardness and low complexity). A puzzling finding in this regard is the fact that neural correlates of accuracy are identified early on in the trial. One possible explanation for this is attentional engagement on the task. If a participant does not actively engage in the task they are more likely to have an incorrect solution. In turn, the likelihood of reaching an incorrect answer due to inattention is higher among instances with low computational difficulty. Together, these patterns would partially explain the marked difference in BOLD activity between correct and incorrect trials in the IPS. However, other alternative explanations are possible. Further work is needed to fully identify the dynamics of effort and attention allocation in computationally complex tasks.

#### Granger causality analysis

PPI analysis provides a description of the functional connectivity (synchronization) between regions based on correlations between simultaneous activity across regions. As such, this analysis is insensitive to temporal directionality in the time series. In contrast, Granger Causality (GC) is defined based on Vector Auto Regression (VAR) models, whereby a vector of ROI signals is driven by a finite number of lags of itself. This allows for gradual excitatory (or inhibitory) impact of one region onto another, that might suggest temporal directionality. This directional effect can be summarized by GC, which emerges when the presence of lags of one variable significantly improves the fit (maximum likelihood value) of another variable. Critical for this study, we expected the underlying neural processes of problem-solving to be internally driven. Specifically, we expected the connectivity patterns to be linked to neural processes whose timing could vary stochastically across trials and participants (e.g., the burst of neural activity does not have to coincide with an experimental intervention such as initial display of items). In order to explore these connectivity patterns we performed a GC analysis on the three ROIs. For this, we ran a VAR model on the ROI time series augmented with the series during the solving stage only, and determined incremental GC of one series on another during problem-solving. This allowed us to estimate effective connectivity GC tests at baseline as well as the GC changes from baseline during the solving stage.

Our analysis shows differential activation between baseline and solving stage. At baseline, during the experiment, we find a bidirectional connectivity between all the ROIs (Fig 14a), which then changes during the solving stage. Specifically, there is a significant change in GC from dACC to rAG. In other words, the dACC (lagged time series) effect on rAG is different between baseline and solving stage. Moreover, we find that during the solving stage there was a significant change in self-activation effect in the dACC and rAG; that is, the lagged time series of each of these two ROIs Granger-cause themselves differentially during the solving stage compared to baseline (Fig 14b).

**Figure 14:**
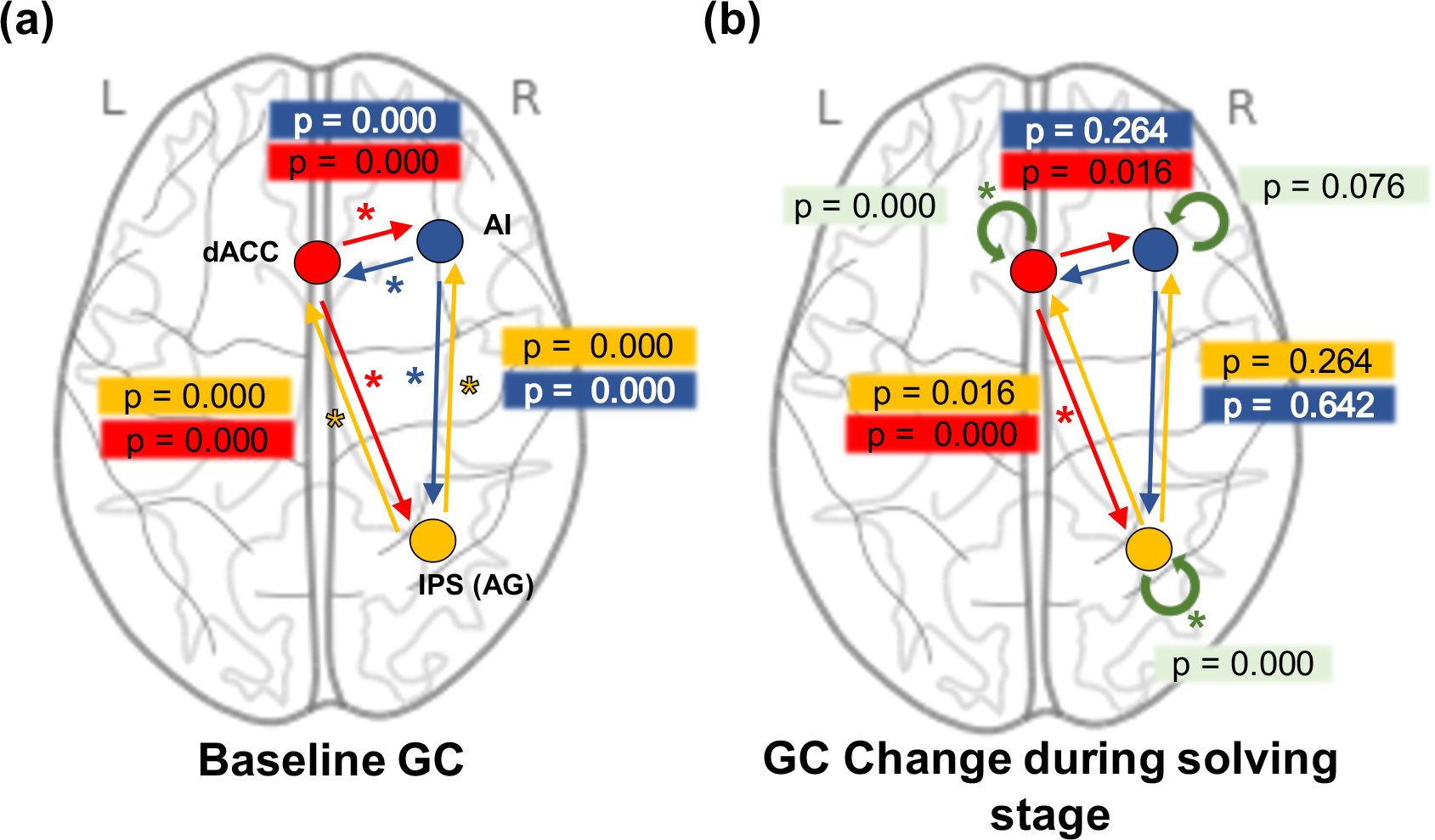
Granger causality results. Effective connectivity estimated via Granger causality between each of three ROIs: dACC, rAI and rAG. **(a)** Represents the baseline connectivity between the regions. **(b)** Represents the changes in effective connectivity during the solving stage compared to baseline. Only three effects survive multiple comparisons correction: An increased connectivity from dACC to rAG and a higher self-modulatory effect on both dACC and AG. *P-values correspond to the GC test uncorrected for multiple comparisons. Asterisks represent significant GC effects FWE-corrected at significance threshold of 0.05*.

Employing the same methodology, we explored whether the change in effective connectivity between solving stage and baseline is modulated by complexity and proof hardness. Specifically, we explored the effects of TCC and satisfiability on GC. We did not find any significant changes in the effective connectivity between high and low TCC instances (all uncorrected p-val*>*0.264) nor between unsatisfiable and satisfiable instances (all uncorrected p-val*>*0.075).

Overall, we found that there was a change in effective connectivity from dACC to IPS during the solving stage of the task. These results extend those previously found in perceptual tasks (Aben et al., 2020), in which regions relevant for the task at hand showed a higher functional connectivity to the dACC during the task. This result further supports previous research that assigns to the dACC a central role in the allocation of control (Shenhav et al., 2013; Dosenbach et al., 2006; Silvetti et al., 2018; Vassena et al., 2017; Holroyd and Yeung, 2012; Alexander and Brown, 2011; Sestieri et al., 2014; Aben et al., 2020; Crottaz-Herbette and Menon, 2006). Interestingly, we did not find a significant change in the effective functional connectivity between rAI and rIPS during the solving stage. These finding match previous research that support a dissociation between dACC and AI (Han et al., 2019; Nelson et al., 2010; Menon and Uddin, 2010; Wu et al., 2019). However, this results seems to be contrary to that found by Sestieri et al. (2014) who found increased functional connectivity between AI and task relevant regions in perceptual and episodic memory tasks. Several possible explanations could be put forward to account for this discrepancy. For instance, the nature of the functional connectivity between rAI and task relevant regions might be task-specific. Specifically, it has been suggested that AI is predominantly involved in processing of internal visceral and motivational information involved in autonomic behavior (Nelson et al., 2010). This type of processing might be more relevant in perceptual and episodic memory tasks compared to the knapsack task, in which mathematical calculations might be more pertinent. Alternatively, other possible explanations for the lack of significant effective connectivity between rAI and rIPS include the lack of statistical power in this analysis, as well as discrepancies in the ROI definition.

Another significant aspect of effective connectivity considered was its link to intrinsic properties of the problem at hand. Our results suggest that the effective connectivity pattern was impervious to the level of computational demand and satisfiability. Of particular relevance, we found that the effective connectivity between dACC and IPS was not modulated by TCC. This suggests that the effect of TCC on control, if any, occurs by generating differential levels of activity within the regions of interest and not via modulation of the functional connectivity between these regions. It is worth noting that this failure to reject the null hypothesis could be due to a lack of power or the exclusion of relevant ROIs from the analysis. We leave it to future research to explore how whole brain connectivity patterns are affected by computational demand.

## Footnotes

3 Significance is determined as follows: order *p* values from small (*k* = 1) to large (*k* = 15); the *k*th test value is deemed to be significant at the level *α* if *p*(*k*) *≤ α/*(*m* + 1 *− k*) where *m* is the number of hypotheses to be tested; here: *m* = 15. If *α* = 0.05 then the smallest *p* should be *≈* 0.0033 for the corresponding test (i.e., the test with smallest *p* value) to reject.

